# 3D-MAESTRO: A scalable, modular, portable pipeline for automated processing of large-scale volumetric brain microscopy data

**DOI:** 10.64898/2026.09.09.750494

**Authors:** Camilo Laiton, Nicholas Lusk, Yoni Browning, Mathew T. Summers, Michael Taormina, Di Wang, Joshua H. Siegle, Holly Myers, Bowen Tan, Polina Kosillo, Erica Peterson, Daphne Toglia, Anna Lakunina, Sasha Burckhardt, Mekhla Kapoor, Tim Wang, John Rohde, Galen Lynch, Shenqin Yao, Sujatha Narayan, Marcus Hooper, Sharon W. Way, Jack Waters, Bosiljka Tasic, Jayaram Chandrashekar, Adam Glaser, David Feng, Sharmishtaa Seshamani, Karel Svoboda

## Abstract

Light microscopy is routinely used to explore the cellular and molecular structure of tissues, but the scale and complexity of data remains a bottleneck for discovery. We introduce 3D-MAESTRO (**3D-M**icroscopy **A**utomation and **E**xecution with **S**calable **T**ools, **R**endering, and **O**rchestration): an automated image processing workflow for large-scale microscopy data, built for scalable execution across cloud and local computing environments. 3D-MAESTRO orchestrates denoising, stitching of image tiles, atlas registration, and segmentation. Its modular architecture permits the integration and benchmarking of new packages, ensuring that performance evolves as more efficient or accurate algorithms emerge. We introduce a new 3D image template for automated registration of mouse brains cleared with aqueous reagents and an efficient method for detection of fluorescent cells. We apply 3D-MAESTRO to lightsheet images of whole mouse brains in the context of diverse anatomical and functional experiments, illustrating high-throughput and reproducible mapping of microscopic structures across the brain.

## Introduction

Brain tissue is organized over spatial scales of nanometers to centimeters (Luo et al. 2018). Understanding the organization of neurons and their circuits requires probing the tissue architecture over these scales simultaneously. Fluorescence microscopy is ideally suited to analyze the structure of biological tissues with molecular contrast and multiplexing. Advances in tissue clearing methods (Spalteholz 1914; Dodt et al. 2007; Hama et al. 2011; Becker et al. 2012; Ertürk et al. 2012; Chung & Deisseroth 2013; Ke et al. 2013; Susaki et al. 2014; Tainaka et al. 2014; Yang et al. 2014; Renier et al. 2014; Hou et al. 2015; Costantini et al. 2015; F. Chen et al. 2015; Chozinski et al. 2016; Sung et al. 2016; Ku et al. 2016) now allow diffraction-limited imaging up to centimeters deep in tissue (Glaser et al. 2025). Combined with turn-key selective plain illumination microscopes (SPIMs) (Stelzer et al. 2021), these methods have made fluorescence microscopy across centimeter-scale tissue volumes, including whole mouse brains (Glaser et al. 2025), routine.

However, challenges in scalability limit the ability of scientists to fully capitalize on high-throughput volumetric imaging. Terabyte-scale datasets must undergo multiple computationally demanding processing stages. These include correction of raw images for optical and illumination artifacts, stitching of multiple image stacks with sub-voxel precision to yield coherent 3D volumes, and registration of these volumes to a standardized coordinate system. Machine-learning models are often essential for detecting and segmenting specific biological features in images. Furthermore, visualizing microscopy data and validating associated data products becomes challenging for terabyte-scale acquisitions.

These sequential steps are burdensome, especially when they exceed the capabilities of standard workstations. For instance, while machine-learning methods are essential for precise structural segmentation and the detection of objects like cells and neurites (Tyson et al. 2021; Attarpour et al. 2025; Friedmann et al. 2020), their reliance on GPU acceleration amplifies computational overhead. Furthermore, integrating results across different specimens, laboratories, and data modalities (such as detecting cells, proteins, or mRNA) requires accurate registration to an atlas (Q. Wang et al. 2020), which demands resource-intensive optimization. Consequently, high-performance or distributed computing has become a prerequisite for completing even routine image analyses within a practical time frame.

Beyond computational cost, these complexities create substantial barriers to scientific reproducibility. Current processing workflows frequently rely on a patchwork of heterogeneous codebases, bespoke scripts, incompatible data formats, and machine-specific configurations. This fragmentation makes pipelines difficult to share and replicate across different computing environments. As a result, data provenance in otherwise well-designed studies is often opaque, undermining the reliability of the findings. Ultimately, the lack of reusable, end-to-end image processing tools remains a primary bottleneck hindering the widespread adoption of high-throughput molecular imaging in intact tissues. Existing tools have often focused on algorithmic innovation for specific tasks rather than delivering end-to-end pipelines. For instance, NuMorph (Krupa et al. 2021) focuses on high-accuracy detection of cellular nuclei. NEATmap (Zheng et al. 2024) is a deep-learning based c-Fos segmentation tool. HiDiver (Johnson et al. 2023) focuses on techniques for registering SPIM and the Allen Mouse Common Coordinate Framework (CCFv3.1) to Magnetic Resonance Histology (MRH) for geometrically correct maps. The BrainGlobe ecosystem (Claudi et al. 2020; Tyson et al. 2021; Claudi et al. 2021; Tyson et al. 2022) provides modular, interoperable tools (e.g., cellfinder, brainrender), but not a reproducible workflow that guarantees results across computational environments. Previous end-to-end pipelines, such as ClearMap (Renier et al. 2016) and the NuMorph toolset (Krupa et al. 2021), do not support distributed computing, and therefore do not scale to terabyte datasets.

We introduce a scalable, automated, and modular pipeline for end-to-end processing of large-scale microscopy datasets (**3D-MAESTRO: 3D-M**icroscopy **A**utomation and **E**xecution with **S**calable **T**ools, **R**endering, and **O**rchestration) (**Figure 1**) (**Table 1**). 3D-MAESTRO also features built-in support for browser-based visualization for all data products via Neuroglancer (https://neuroglancer-demo.appspot.com). We developed specific implementations optimized for images of whole mouse brains, but 3D-MAESTRO can be used for general 3D microscopy platforms and applications with minor changes. Our design emphasizes:

- scalability through parallel and distributed computation,
- standardization based on FAIR-aligned (Wilkinson et al. 2016) open data formats,
- modularity enabled by containerization and integration with a workflow engine, and
- portability across cloud computing environments, on-premises HPC systems, and local workstations.

The 3D-MAESTRO architecture allows emerging algorithms to be incorporated seamlessly as they become available. We illustrate the pipeline’s versatility with three types of neuroscience applications: characterizing tools for targeting specific cell types for gene expression, such as adeno-associated virus (AAV) enhancer reagents and transgenic mouse lines; brain-wide mapping of connectivity by detecting cells and neurites in multicolor circuit tracing experiments; and localizing measurements of neural function including cFOS activation and *in vivo* electrophysiological recordings. We demonstrate that our pipeline transforms large-scale SPIM microscopy images into reproducible, quantitative data while delivering the scalability and reliability necessary to support future advances in imaging, algorithms, and computation.

**Table 1.**
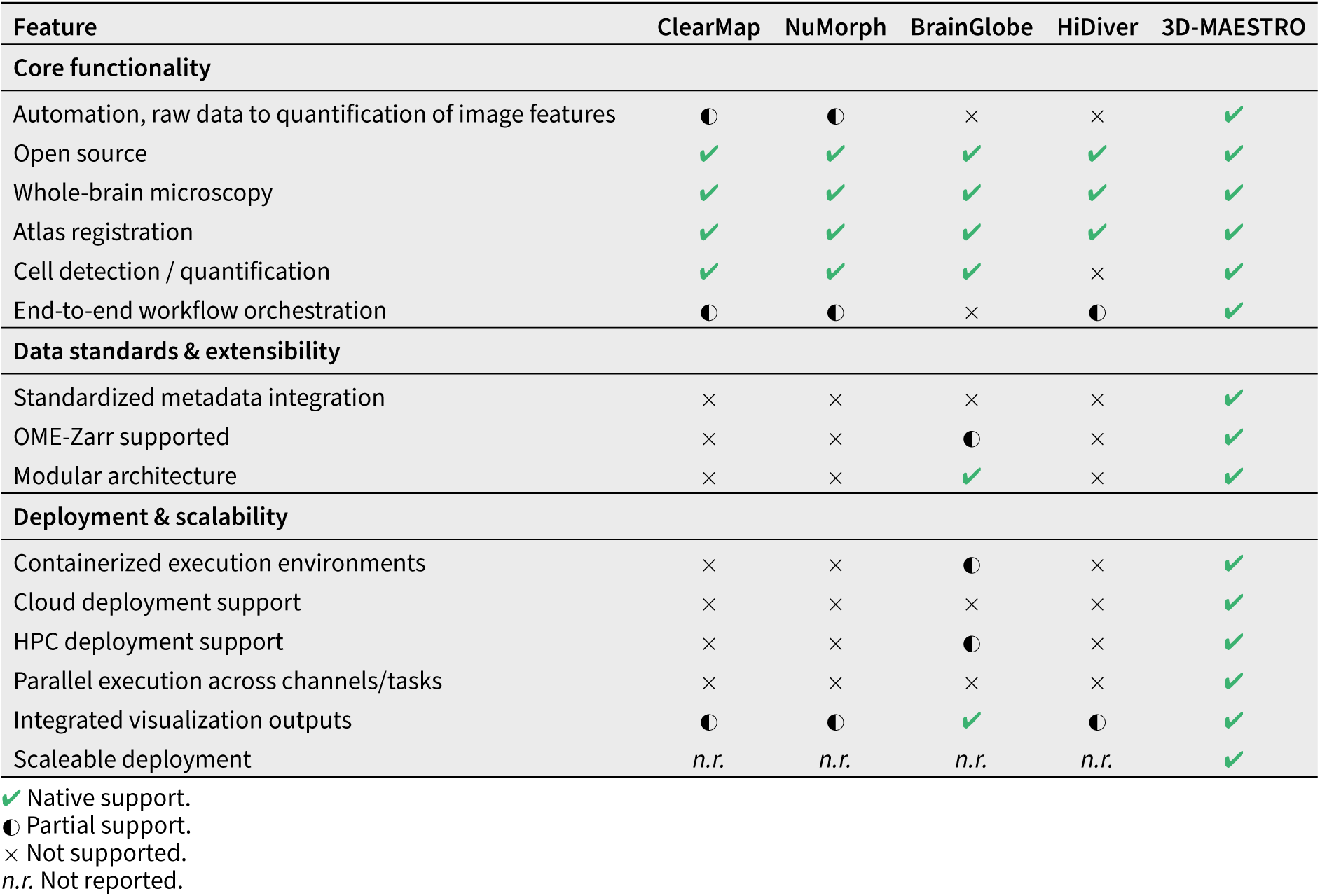
Comparison of representative tools for whole-brain microscopy data processing.

## Results

### 3D-MAESTRO: an end-to-end processing pipeline for large-scale microscopy data

3D-MAESTRO orchestrates multiple stages of microscopy data processing (**Figure 1**). The core stages produce a multiscale volume, which enables browser-friendly hierarchical visualization via automatically generated Neuroglancer views. To create multiscale image volumes, 3D-MAESTRO first performs data ingest and metadata preparation, followed by image artifact correction, image tile alignment, and image volume fusion (**Figure 1**). Additional application-specific workflows such as atlas registration and cellular feature detection aids the transformation of large-scale microscopy datasets into biologically meaningful insights. Neuroglancer-based visualization enable rapid quality control and validation of data products. Every stage of 3D-MAESTRO creates standardized outputs, allowing modules to be readily replaced. Four design principles underpin the implementation of 3D-MAESTRO: scalability, standardization, modularity, and portability.

#### Scalability

To enable scalable processing, we design each stage for parallel or distributed processing and ensure that resources are matched to the needs of each step. Given that individual datasets can range from hundreds of gigabytes to several terabytes, this is a critical feature. We use Spark (Zaharia et al. 2010) (Hörl et al. 2019), which allows datasets to be processed in smaller chunks concurrently. For cell quantification, we employ Ray (Moritz et al. 2018), a distributed task framework well suited to independently assigning detected cells to atlas regions.. Cell detection steps are optimized on GPUs using CuPy (Okuta et al. 2017), which accelerates the array-based computations underlying image processing and inference. Across modules, we rely on Python multiprocessing and sharedmemory strategies to handle large scale data (from local and cloud locations). Additionally, we leverage distributed computing to parallelize processing across the fluorescence channels present in each dataset. Together, these strategies enable the pipeline to scale with dataset size and allow users to flexibly allocate computing resources depending on their data and analysis needs. Over the past two years, we have processed approximately 4,000 whole-brain datasets using both cloud and on-premises HPC resources.

**Figure 1.**
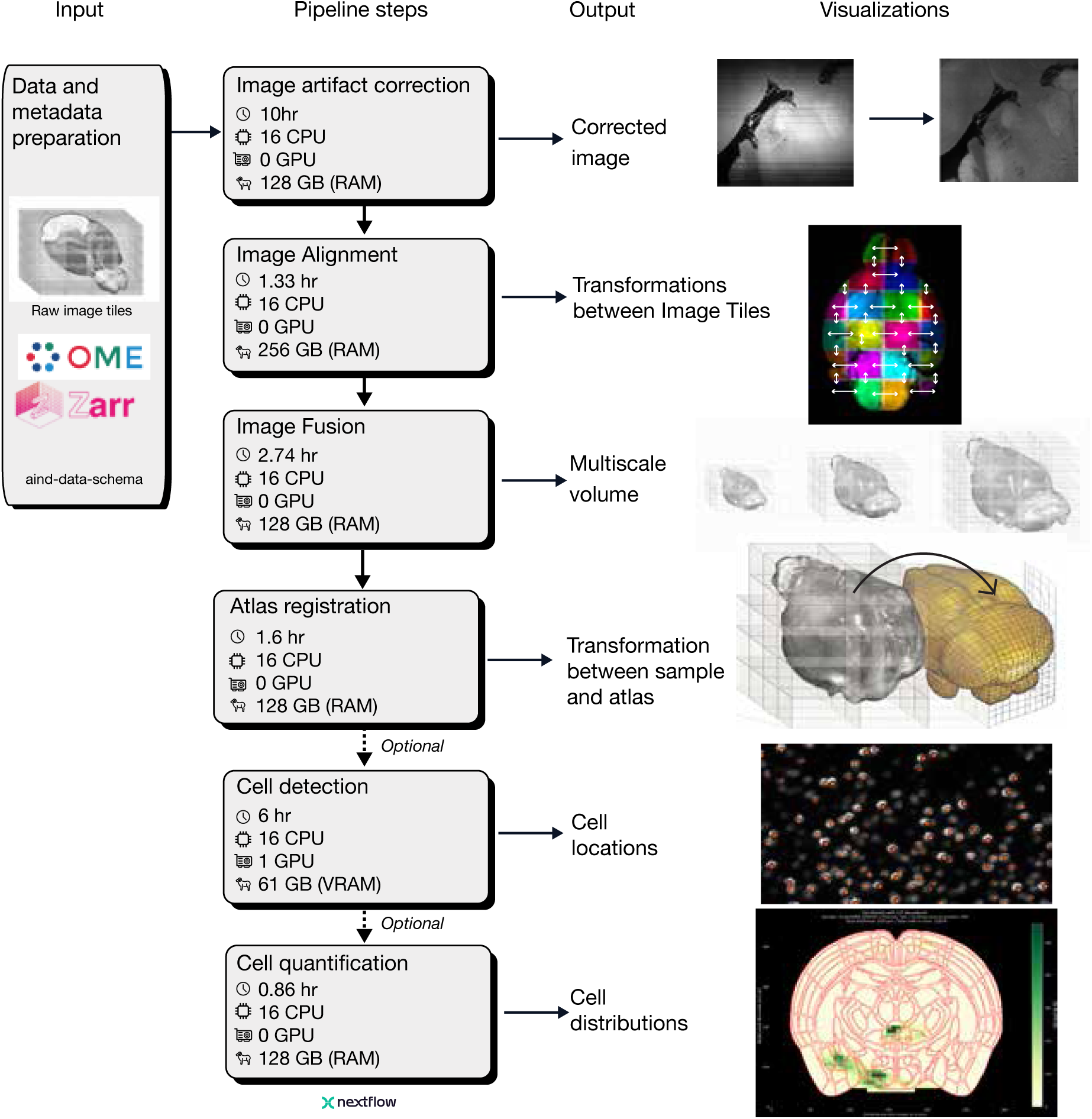
3D-MAESTRO overview. Flow diagram of the image processing pipeline applied to SPIM data and cell localization in whole mouse brains. Average resources and real time for processing one whole mouse brain dataset (3 channels, acquired at 1.8um x 1.8um x 2um - totally 1TB) are reported in each box. **Input**, images in standard formats and metadata are imported for processing. **Image artifact correction**, enhance image quality (Figure 2). **Image alignment**, align image stacks for near pixel perfect overlap at tile boundaries. **Image fusion**, merge image stacks into a single, multiscale volume (Figure 2). Beyond these core stages, additional workflows are required to map labeled cells across the mouse brain. **Atlas registration**, register volume to a standard coordinate system for the mouse brain (Figure 3). This step is critical for combining data across multiple experiments and link the data to other maps (e.g. gene expression). **Cell detection**, find centroids of labeled cells (Figure 4). **Cell quantification (see Methods)**, common example application, counting cells in brain volumes. Dashed arrows indicate optional analyses, to be replaced with other analysis modules as needed. The images on the right illustrate processing steps. Outputs of every step can be visualized with Neuroglancer. https://neuroglancer-demo.appspot.com/

#### Standardization

3D-MAESTRO is built around community standards. For 3D volumes, we use the OME-Zarr file format (Moore et al. 2023), a chunked, multiscale format designed for cloud-native workflows. Because volumes are stored as independently addressable chunks across multiple resolution levels, OME-Zarr enables fast, parallel, on-demand streaming directly from cloud object storage. This allows users to access arbitrary sub-volumes without downloading entire datasets. This chunked, multiscale organization also makes the data immediately compatible with Neuroglancer, a browser-based rendering tool for volumetric data that requests only the chunks and resolution levels needed for the current view, supporting interactive exploration of terabyte-scale volumes on standard hardware without local copies of the data. Alongside the imaging data, all metadata are captured using the AIND data schema(https://github.com/AllenNeuralDynamics/aind-data-schema), an open-source schema that provides a consistent structure for microscopy metadata and detailed system logs for imaging parameters.

#### Modularity

To facilitate applications across diverse data sets analyzed with multiple algorithms, 3D-MAESTRO features a plug-and-play architecture driven by two design choices. First, we use Nextflow (2025), a system for reproducible scientific data processing. A Nextflow script composes independent processes into increasingly complex pipelines, with each process executing a given tool or scripting language. By specifying the process inputs and outputs, Nextflow coordinates the execution of tasks, manages data transfer between them, enables parallelization and ensures transparent and reproducible input-output relationships. Nextflow supports modularity by treating each process as a self-contained, reusable building block with clearly defined inputs, outputs, and execution environments. The individual module logic is decoupled from the overall workflow structure making it easy to rearrange, replace, or extend pipeline components, such as swapping algorithms or incorporating novel pre- and post-processing steps— without modifying the core workflow. Second, we ensure that inputs and outputs for each component follow a consistent schema (OME-Zarr for images and metadata consistent with aind-data-schema) as described in the standardization section. These two design choices together enable plug-and-play interchangeability within the pipeline. For example, it is straightforward to compare different algorithms and update specific processing steps as new algorithms become available.

#### Portability

3D-MAESTRO runs on diverse compute backends. The pipeline was built cloud-first using Code Ocean (Cheifet 2021), a platform for reproducible scientific computing. Code Ocean natively supports NextFlow pipelines run on scalable cloud resources. Because NextFlow’s open-source engine supports a variety of backends, the pipelines also run on local workstations and HPC Slurm clusters. All source code is hosted publicly on GitHub, with public containers for each process published in GitHub Container Registry.

### Image processing

#### Image artifact correction

SPIM suffers from two types of artifacts that impair image quality, which can hinder accurate analysis. Correcting these artifacts facilitates downstream image processing and more reliable quantification, particularly in measurements sensitive to intensity variations across samples (e.g., segmenting cellular structures based on their brightness).

##### Vignetting

A variety of mechanisms produce repeated, predictable variation in image brightness (**Figure 2A**). Vignetting occurs due to angular sensitivity of detectors, reduced light collection efficiency of the lenses or imperfect optical calibration of the microscope (Goldman 2010; Tomazevic et al. 2002; Piccinini et al. 2012). Similar variations in image brightness are caused by non-uniform illumination and enhanced photobleaching in image regions with overlap of image tiles. We apply retrospective correction of vignetting (Tomazevic et al. 2002) using BaSiC (Peng et al. 2017), which estimates the function required to correct brightness variations in microscopy images. We evaluate the correction by measuring the line difference spread (LDS) and pixel value spread (PVS) between corrected and uncorrected images (see Methods).

**Figure 2.**
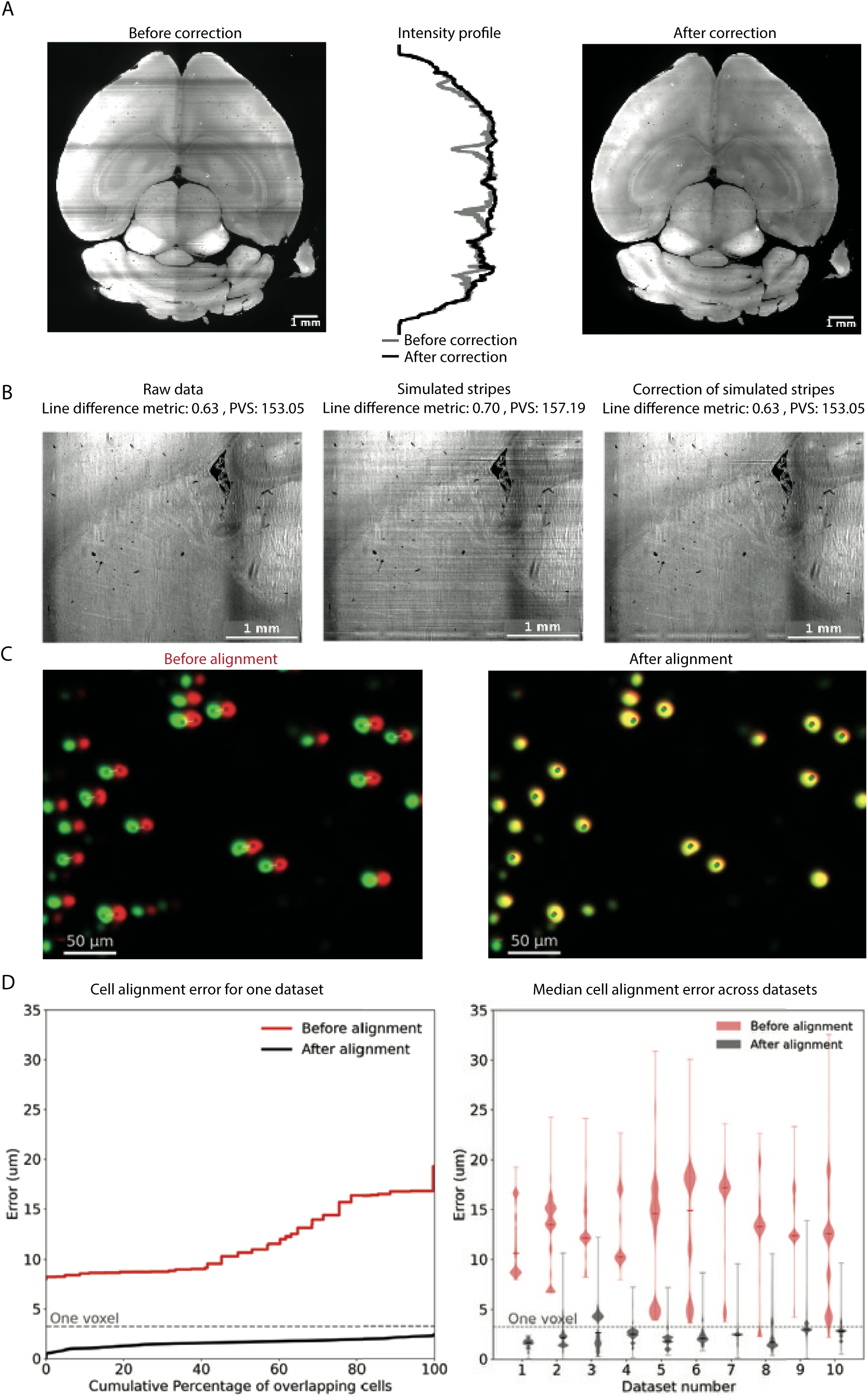
Image Processing. **A.** Raw (uncorrected, left) and corrected (right) images of a whole brain volume. Center, intensity profile before (grey) and after (black) correction **B.** Images show a raw image (left), same image with added stripe artifacts (middle), and image after applying the destriping algorithm. The line difference metric is a measure of striping but depends on image content and thus does not have an absolute meaning. **C.** Locations of cells with fluorescent nuclei before and after alignment in an overlap region between two tiles (red and green). Overlapping cells appear yellow. Double detections before alignment are corrected with alignment **D.** Inter-Image Cell Correspondence Error (ICCE) for cells within overlapping regions before (red) and after (black) alignment. Left, ICCE error as a function of the cumulative percentage of cells in a representative dataset. Right, distributions of ICCE error for ten datasets before (red) and after (black) stitching (violin plots with medians). Dataset 1 corresponds to the dataset shown in the left panel.

##### Striping

Striping artifacts arise due to the absorption and scattering of the light sheet by inhomogeneities within the tissue, such as blood, melanin, air bubbles, or subcellular structures (Ricci et al. 2022; Jacques 2013). Striping artifacts are typically smaller than vignetting. In SPIM images, striping artifacts usually appear along the direction of light-sheet propagation, which is perpendicular to the detection axis, creating horizontal stripes. Retrospective stripe artifact removal is typically addressed through optical or digital filtering (Ricci et al. 2022) using image processing algorithms to correct data (Liang et al. 2016).

We adapt the open-source software pystripe (Kirst et al. 2020), which suppresses stripe artifacts by decomposing the image into wavelet subbands and damping the stripe-associated frequency components via FFT filtering. This filtering is applied independently to the image’s foreground and background regions, each with its own filter bandwidth, to account for their differing levels of stripe contamination. In addition, we have extended pystripe’s capabilities to support the OME-Zarr format and read data from cloud-storage services.

To assess the effectiveness of destriping, we compare LDS and PVS (Methods) before and after artifact removal. We tested our method based on stripes added into real data (Methods) (**Figure 2B**). In addition, stripe removal was evaluated by manual inspection.

#### Stitching

When imaging specimens that exceed the microscope field of view, volumes are acquired as overlapping image stacks to cover the entire 3D sample. These tiles require stitching to generate a multiscale volume for downstream analysis. Stitching comprises two computational steps: image alignment and image fusion. Image alignment calculates the optimal spatial relationships between tiles. Image fusion blends these tiles into a single multiscale volume.

Although both steps are computationally intensive, existing software packages such as TeraStitcher (Bria & Iannello 2012) and BigStitcher (Hörl et al. 2019) are designed to address these demands. We leveraged the modular architecture of 3D-MAESTRO to benchmark the two tools across 1,473 datasets using EC2 R4.8xlarge instances (32 virtual cores and 244 GB RAM). Note that this was used for the benchmarking experiment which is different from Table 4, which reports what the routine pipeline runs. BigStitcher achieved approximately twice the processing speed of TeraStitcher (Supplementary Figure S1), despite performing the more computationally demanding task of full 3D alignment, whereas TeraStitcher uses 2D maximum-intensity projections for alignment. BigStitcher was also more stable, and more efficient than TeraStitcher for image fusion. Based on these results, we selected BigStitcher as the stitching framework for 3D-MAESTRO.

We next assessed alignment quality on diverse images used for quantifying labeled cells across the brain. The Inter-Image Cell Correspondence Error (ICCE;see Methods) measures the distance between corresponding cells that appear in overlap regions (**Figure 2C**). For aligned images ICCE should be near zero. The median ICCE across datasets decreased from 12.6µ*m* (range, 7.4 – 17.2µ*m*) before alignment to 2.2µ*m* (range, 1.6 – 3.0µ*m*) after alignment, a 5.7-fold reduction (**Figure 2D**). Since the 3D diagonal image voxel size is 3.24um, the image alignment error is on the order of one-voxel. Fused volumes showed no duplication of cells within overlap regions. This precision demonstrates the robustness and reliability of the alignment method across a wide range of imaging datasets.

### Registration to a common coordinate system

It is often of interest to integrate image data across individual subjects. This can be accomplished by registering individual experiments to a common reference space (or coordinate system), by warping the imaged volume to a template brain volume (e.g. the CCFv3.1 template(Q. Wang et al. 2020)). For many applications, registration must be commensurate with the tissue’s finest scale of organization, such as laminar features and brain nuclei in rodent brains, which can be smaller than 100µm. Precise registration to a common reference space also provides a multi-modal context to any measurement. For example, anatomical measurements can be interpreted in the context of molecular taxonomies (Yao et al. 2023) and functional data, such as electrophysiology (Liu et al. 2021; S. Chen et al. 2024; Bennett et al. 2024). High-resolution registration of measurements across subjects, laboratories, and modalities is required for an integrated understanding of organismal biology.

Several open-source tools promise registration between brain volumes (Q. Wang et al. 2020; Fischl 2012; Klein et al. 2010; Tustison et al. 2021). However, the performance of these algorithms relies on intensity-based similarity metrics and performance degrades severely when applied across images acquired with different types of microscopes or contrast mechanisms (Perens et al. 2021). These challenges are evident when comparing SPIM images and serial 2-photon tomography images used to generate the Allen Common Coordinate Framework (CCFv3.1) anatomical template (STPT) (**Figure 3 A-G**) (Q. Wang et al. 2020). Notably, white matter tracts appear light in SPIM because of high autofluorecence, but dark in STPT because of strong light scattering (**Figure 3 C,F**; quantification in **Figure S2B,C**).

Two strategies have been traditionally used to automate alignment across contrast mechanisms. The first strategy uses segmentation-guided registration, where anatomical regions are segmented and mapped between modalities to guide alignment (Qiu et al. 2024; Roston et al. 2025). This approach typically requires extensive 3D manual annotation on individual subjects. Moreover, segmented region boundaries may not easily be matched across different contrast mechanisms, causing systematic warping errors. The second strategy uses an image template specific to each contrast mechanism, commonly generated by averaging multiple samples of a given modality (Perens et al. 2021; Pisano et al. 2022; Liu et al. 2021). This method allows registering new samples by first aligning to the modality-specific template (an intra-modal task), which itself has a mapping to the cross-modal atlas defined *a priori*. The laborious manual annotation required for registration across contrast mechanisms needs only occur once, for the modality-specific template.

We constructed a new template brain volume from SPIM images of aqueous cleared tissue (Liu et al. 2021). We first imaged far-red (639 nm excitation, 660–680 nm emission) autofluorescence signal from ten whole mouse brains. Images were standardized across samples using bias-field correction (Tustison et al. 2010) and intensity normalization (**Figure S3A**). We then constructed a high-contrast, high-resolution (10 µm) SPIM template (ST) by warping brain volumes to each other and averaging (**Figure 3G**, **Figure S3**). Fine details including barrel fields in somatosensory cortex (**Figure 3H**) as well as barrelettes and barreloids in the medulla and thalamus (**Figure S3E-H**) are clearly visible in the ST (Q. Wang et al. 2020). To create a mapping between the ST and the STPT related to CCFv3.1, we combined intensity-based gradient metrics with segmentation-guided alignment (**Figure 3I**). Annotations supporting segmentation-guided alignment came from small (e.g. LGd, act) and large brain regions (e.g. cortex, cerebellum). This workflow enables automated cross-modal registration of new samples by first warping individual brain volumes intra-modally to the ST, and then leveraging the re-usable mapping of the ST to CCFv3.1 (**Figure 3J**).

Template generation and registration were implemented using Advanced Normalization Tools (ANTs) toolkit (**Figure 3H**) (Tustison et al. 2021; B. Avants et al. 2009; B. B. Avants et al. 2011). The modularity of 3D-MAESTRO ensures that as new atlases or templates become available, they can be easily integrated. For instance, additional structural annotations can be incorporated to further refine registration if local inaccuracies are identified. Moreover, because the ANTs-derived template-to-atlas transformations are one-to-one, different mappings can be swapped or evaluated directly - without the computational cost of re-registering each individual brain dataset. This design provides both flexibility and scalability as methods and datasets continue to evolve.

**Figure 3.**
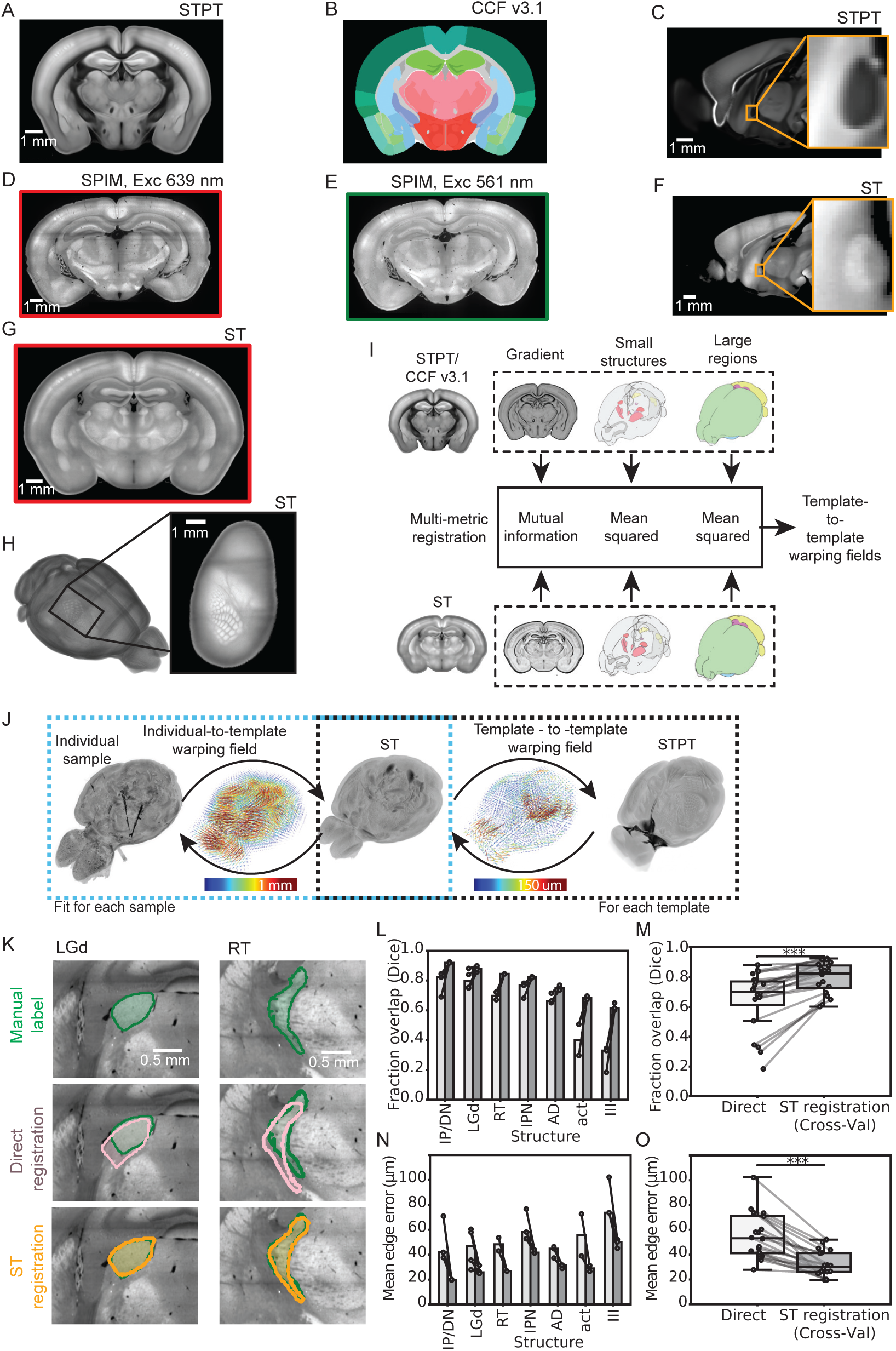
Registration to a common coordinate system. **A.** Slice from the serial 2-photon tomography template (STPT) defining the mouse Common Coordinate Framework (CCFv3.1). **B.** Annotations (CCFv3.1.1). **C.** Highlight of the anterior commissure in the STPT. **D.** Slice through an individual mouse brain imaged using SPIM (red channel, excitation 639 nm/emission 667 nm). **E.** Green channel (excitation 561 nm/ emission 593 nm). **F.** Same as C, but for SPIM images. Note the qualitative difference in contrast with C. **G** Slice of the SPIM template (ST). **H.** 3D ST; inset, barrel field in autofluorescence. **I.** Process for registering the (ST) to the STPT. **J.** Registering an individual SPIM image volume to ST and then the STPT. Transforms are shown as vector fields, with only the non-linear component shown. **K.** Comparing registration methods. Top, manual annotation of LGd (right) and RT (left). Middle, direct-to-STPT registration. Bottom, our ST-based registration pipeline. **L.** Dice scores comparing registered labels to manual annotation for direct registration on the stitched image (light gray) and ST-based registration (dark gray). **M.** Same as in *L* but shown as a population with quartiles. *s indicate rank sum test: ***p<=.001. **N,O.** Mean distance to the closest surface, shown as in **L, M** *LGd*: Dorsal part of the lateral geniculate complex. *IP/DN*:Interposed nucleus/Dentate nucleus (cerebelar nuclei). *RT*:Reticular nucleus of thalamus *AD*:Anterodorsal nucleus *act*:anterior commissure, temporal limb *III*:Oculomotor nucleus.

We next evaluated registration to a standard coordinate system. In mouse brain studies, registration to the CCFv3.1 (Q. Wang et al. 2020) (**Figure 3 A-C**) is the standard for integration with community datasets. The CCFv3.1 includes a multi-resolution STPT template, a related coordinate system, and a set of area annotations on this template (**Figure 3A,B**). We evaluate registration of whole mouse brain SPIM images to this STPT. To create an evaluation dataset, we used five test brains to estimate candidate registrations to the CCFv3.1 Atlas. In each of these brains, a subset of seven test structures (LGd, IP/DN, RT, IPN, AD, act, and III) were manually annotated in 3D. Each brain was then registered to the 25 um STPT template either by direct intensity-based registration (“direct”) or using ST-based registration.

The differences across methods can account for major effects in downstream analyses, especially for small structures (**Figure 3 K**). The ST-based registration improved overlap between structures (Dice scores, **Figure 3L,M**, see also **FigureS2D**) and surface-to-surface distances (**Figure 3N,O**; **Figure S2E**), providing state-of-the art performance (Liu et al. 2021).

### A deep learning algorithm for efficient and accurate cell detection in SPIM images

We use the cell detection plugin, which is optional in 3D-MAESTRO to detect and localize brain cells expressing fluorescent molecules in SPIM data. This is a widely used task in studies of gene expression, neural development, anatomy and other applications. The labeling method can have large effects on signal and background levels, as well as the density and spatial overlap of labeling. Given the diversity of possible imaging and labeling conditions, accurate and robust detection of cells is a challenging problem (**Figure 4A**). Existing algorithms can be divided into instance detection (Tyson et al. 2021; Murakami et al. 2018; Iqbal et al. 2019) and segmentation methods. (Stringer et al. 2021; Hörst et al. 2024; Achard et al. 2024; L.-W. Wang et al. 2023). Instance detection directly identifies the locations of individual objects and assigns object-level labels. It requires only object-level localization, making it computationally efficient, easy to annotate, and well-suited for scaling to large datasets. Segmentation methods predict class labels for every pixel or voxel to delineate object boundaries. Segmentation enables detailed morphological analyses, including measurements of cell size and shape, but requires substantially denser annotations and greater computational complexity. Here, our primary goal is the accurate localization of individual cells allowing for whole-brain quantification. We therefore implemented a two-step instance detection framework with a focus on computational efficiency and scalability.

In the first step, cell locations are proposed using classical, GPU-accelerated image processing (background normalization, blob detection, and shape-based filtering; see Methods). Proposal generation is deliberately parameterized liberally, minimizing false negatives at the cost of admitting spurious detections. In the second step, an 18-layer ResNet classifier (He et al. 2015) separates true cells from false positives. The classifier operates on both the signal and autofluorescence background channels, allowing it to reject broad-spectrum artifacts — such as tissue edges and nonspecific fluorescence — that are missed by algorithms considering only a single channel (see Methods for training details). Both components are modular and can be independently updated or replaced.

**Figure 4.**
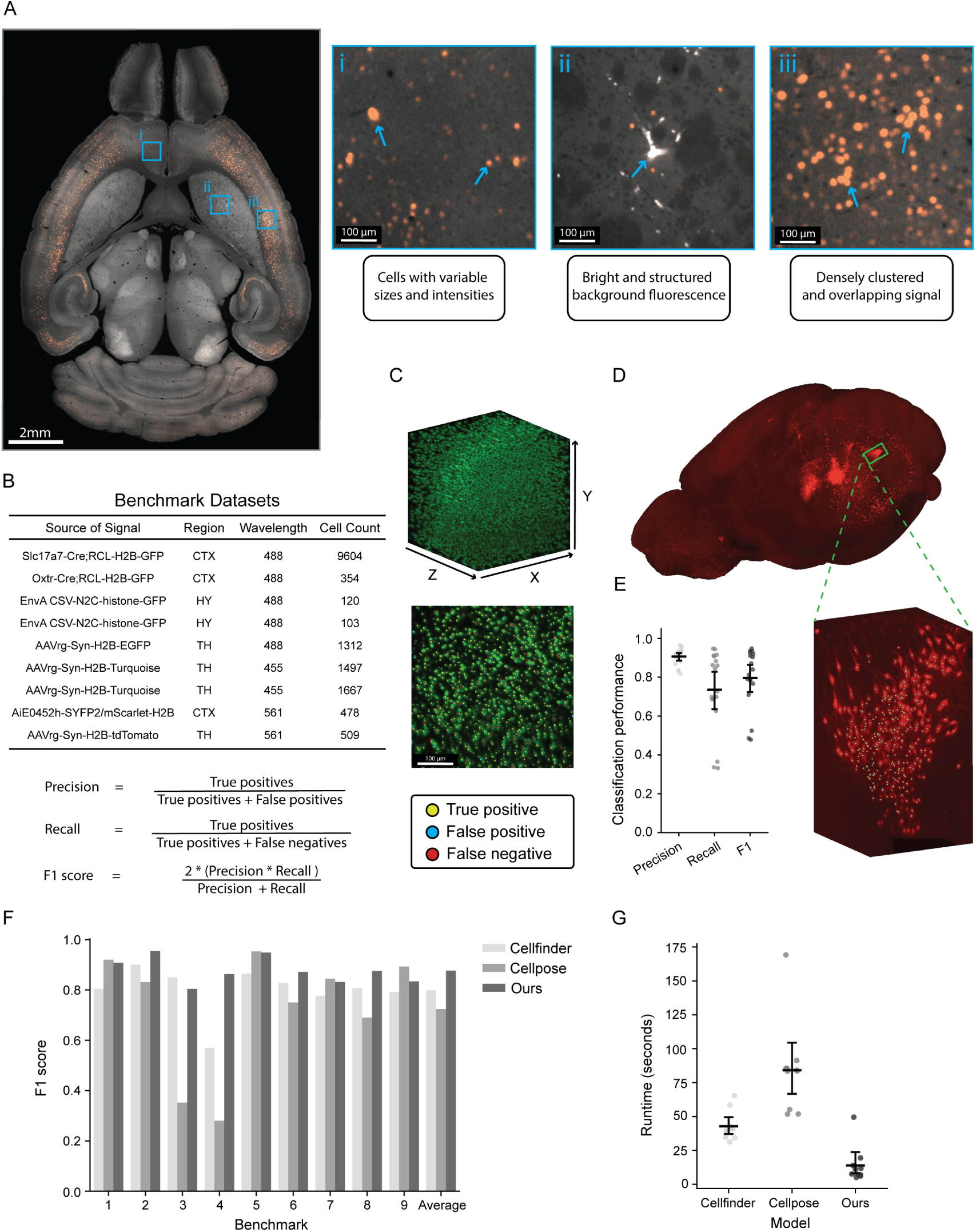
A deep learning pipeline for efficient and accurate cell detection in SPIM images. **A.** Horizontal section through a WT C57BL/6J brain volume with a retro-orbitally injected AiE2333m-SYFP2/mScarlet-H2B (red) and autofluorescence (white). Regions i-iii highlight the diversity of signal across the brain, including cells with high variability in intensity and size, cells intermingled with background fluorescence, and clustered cells with signal overlap. **B.** Benchmark datasets for testing cell detection performance. 9 held out datasets were created, consisting of fully annotated blocks (256 *×* 256 *×* 256 or 256 *×* 256 *×* 128 voxel block; voxel volume, 1.8 *×* 1.8 *×* 2.0 µm^3^). Benchmarks cover a broad spectrum of signal across brain regions, channels, densities. **C.** Classification performance on benchmark volume shown as max intensity projections (top) and a slice through the volume (bottom) with locations identified by our algorithm. Each point is classified as true positives (yellow), false positives (cyan), and false negatives (red) relative to ground truth human annotation (human labelled cells). Overlapping labels are the result of points with similar X and Y coordinates at different depths of the max projection. Cells with no labels have annotations outside the max projection range. **D.** Max projection of whole brain signal from a Dbh-Cre-KI driver line crossed to a Cre+FlpO reporter (AI65) expressing cytosolic tdTomato after injection of AAVretro-DIO-FlpO. **E.** Model performance on 18 datasets expressing cytosolic labeling after fine-tuning on similar data (left) and max projection of classification results of cytosolic expression in the Pons (right) **F.** Performance of multiple algorithms across benchmark datasets. **G**. Processing speed across algorithms on the benchmark dataset sub-volumes. Runtime for two-phase algorithms include both proposal generation and classification phases. Algorithms were tested using 16 CPUs, 1 V100 GPU, and 64Gb RAM.

We tested our algorithm on a benchmark dataset with nine fully-annotated brain volumes (either a 256 *×* 256 *×* 256 or 256 *×* 256 *×* 128 voxel block; voxel volume, 1.8 *×* 1.8 *×* 2.0 µm^3^) containing cells expressing fluorescent proteins in their nuclei (Methods). The nine volumes were selected to cover a variety of expression densities, fluorescence colors and brain regions (Figure 4B). Each volume was manually annotated by an expert annotator. To ensure evaluation integrity, these brain volumes were not used for training the classification model (see Methods).

We computed precision, recall and F1 scores for each benchmark dataset (**Figure 4B, C**). The model prediction may result in a cell being correctly identified as a true positive, but not at the same voxel location as the manual annotation. To account for this, a proximity based algorithm was used to match model predictions to associated benchmark annotation.

For detecting cells labeled with cytosolic fluorescent protein (**Figure 4D**), the model was fine-tuned using 7,500 annotations of cells expressing cytosolic tdTomato. The model was evaluated on eighteen brains from a Dbh-Cre driver line (Tillage et al. 2020) crossed to a Cre+FlpO reporter (AI65) (Madisen et al. 2015) following AAVretro-DIO-FlpO (Zingg et al. 2017) injections. Each dataset was fully annotated within the pons (Su et al. 2026) and evaluation was also limited to the pons, (**Figure 4D-E**). This shows that fine-tuning with only a relatively small number of annotations produced a model with robust performance across the 18 datasets (Precision: 0.91 *±* 0.043, Recall: 0.74 *±* 0.21, F1: 0.80 *±* 0.15.

We compared our detection algorithm with two widely used tools: Cellfinder (cell detection) (Tyson et al. 2021) and CellPose (cell segmentation) (Stringer et al. 2021). As Cellfinder also uses ResNet classification, we use our trained ResNet with their detection algorithm for comparison. The Cellpose model was fine-tuned on in-house datasets to provide a fair comparison. All approaches were assessed for F1 scores and processing speed (16 CPUs, 1 V100 GPU, and 64GB RAM). For evaluation of CellPose, the centroid of each instance was assigned as the detection location. Across the nine benchmark volumes, our model outperformed both Cellpose and Cellfinder in overall F1 score (Cellfinder: .80; Cellpose: 0.72; Ours: 0.88) and runtime (Cellfinder: 42.81*s ±* 11.49*s*; Cellpose: 84.12*s ±* 35.73*s*; Ours: 13.90*s ±* 14.06*s*) (**Figure 4F-G**).

**Figure 5.**
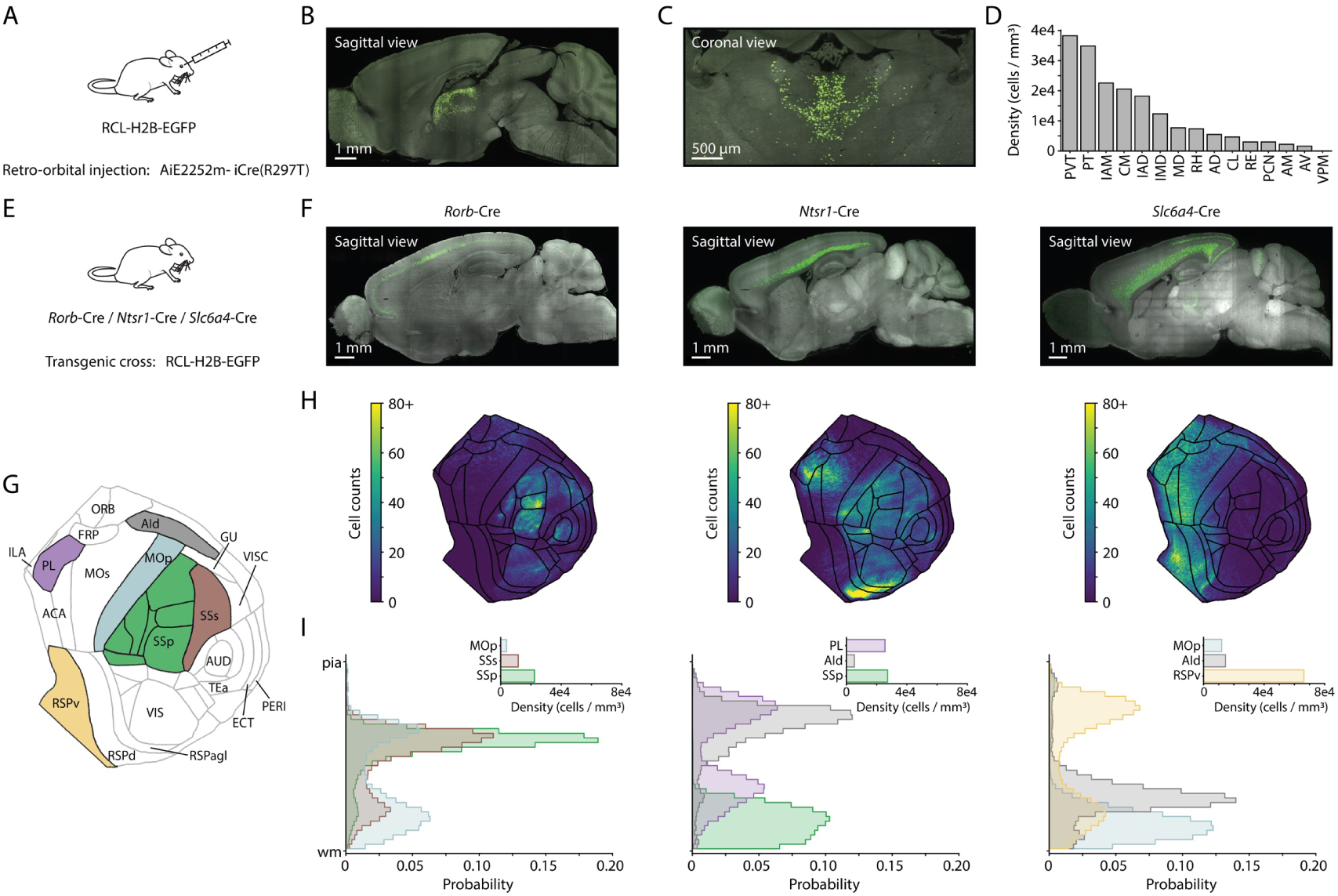
Characterization of tools for cell type-specific gene expression. **A.** Retro-orbital delivery of AiE2252m-iCre(R297T) in a reporter mouse expressing nuclear GFP. **B.** Sagittal view of labeled mouse brain. **C**. Same as **B**, but coronal view of thalamus. **D**. Densities of labeled nuclei across thalamic nuclei. **E**. Crosses of transgenic mice expressing Cre with reporter mouse expressing nuclear GFP. **F**. Sagittal view of mouse brain samples from *Rorb-*Cre (*lek*), *Ntsr1-*Cre (*middle*), or *Slc6a4-*Cre (*right*) crosses. **G**. Cortical flatmap with area labels. **H**. Cortical flatmaps showing the spatial distributions of detected cells. **I.** Detected cells as a function of depth in the cortex, normalized to cortical thickness. Colors correspond to areas highlighted in **G**. ACAd – anterior cingulate area, dorsal part, ACAv – anterior cingulate area, ventral part, AId – agranular insular area, dorsal part, AIp – agranular insular area, posterior part, AIv – agranular insular area, ventral part, AUDd – dorsal auditory area, AUDp – primary auditory area, AUDpo – posterior auditory area, AUDv – ventral auditory area, ECT – ectorhinal area, FRP – frontal pole, GU – gustatory areas, ILA – infralimbic area, MOp – primary motor area, MOs – secondary motor area, ORBl – orbital area, lateral part, ORBm – orbital area, medial part, ORBvl – orbital area, ventrolateral part, PERI – perirhinal area, PL – prelimbic area, RSPagl – retrosplenial area, lateral agranular part, RSPd – retrosplenial area, dorsal part, RSPv – retrosplenial area, ventral part, SSp – primary somatosensory area, SSp-bfd – primary somatosensory area, barrel field, SSp-ll – primary somatosensory area, lower limb, SSp-m – primary somatosensory area, mouth, SSp-n – primary somatosensory area, nose, SSp-tr – primary somatosensory area, trunk, SSp-ul – primary somatosensory area, upper limb, SSp-un – primary somatosensory area, unassigned, SSs – supplemental somatosensory area, TEa – temporal association area, VISa – anterior visual area, VISal – anterolateral visual area, VISam – anteromedial visual area, VISC – visceral area, VISl – lateral visual area, VISli – laterointermediate visual area, VISp – primary visual area, VISpl – posterolateral visual area, VISpm – posteromedial visual area, VISpor – postrhinal area, VISrl – rostrolateral visual area, AD – anterodorsal nucleus, AM – anteromedial nucleus, AV – anteroventral nucleus, CL – central lateral nucleus, CM – central medial nucleus, IAD – interanterodorsal nucleus, IAM – interanteromedial nucleus, IMD – intermediodorsal nucleus, MD – mediodorsal nucleus, PCN – paracentral nucleus, PT – parataenial nucleus, PVT – paraventricular nucleus of the thalamus, RE – reuniens nucleus, RH – rhomboid nucleus, VPM – ventral posteromedial nucleus.

### Characterization of tools for cell type-specific gene expression

We next showcase 3D-MAESTRO’s capabilities in the context of several types of experiment commonly performed in neuroscience. Genetic tools such as transgenic mouse lines (Gong et al. 2003; Gong et al. 2007; Harris et al. 2014) and enhancer AAVs (eAAVs) (Dimidschstein et al. 2016; Hrvatin et al. 2019; Graybuck et al. 2021; Ben-Simon et al. 2025) are foundational tools for accessing genetically defined cell types for gene expression. Determining the completeness and specificity of labeling, as well as possible off-target expression, are important steps in developing and characterizing candidate genetic tools. This requires mapping brain-wide expression patterns at high throughput. We combined transgenic mice and eAAVs expressing Cre with reporter mice to express nucleus-localized GFP. 3D-MAESTRO was used to map GFP-expressing cells across the brain.

We first characterized retro-orbitally delivered AiE2252m-iCre(R297T), an eAAV designed to target expression to the dorsal midline thalamus (including paraventricular, PVT and PT, respectively) (**Figure 5A**). We predominantly observed labeled cells in thalamus, with a small number of labeled cells in hypothalamus and medulla (**Figure 5B, C**). Within thalamus, the density of labeled cells was highest in PVT and PT (**Figure 5D**). Additional labeling was observed in other midline thalamus structures such as the intermediodorsal (IMD) and ventral midline nuclei, as well as intralaminar nuclei. We did not observe labeling in sensory structures such as the ventral posteromedial nucleus (VPM), reflecting the distinct transcriptomic profiles of these cells (Yao et al. 2023; Turner et al. 2025).

We next used 3D-MAESTRO to characterize the expression of three transgenic mouse lines targeting excitatory neurons in specific cortical layers (Harris et al. 2014; Gong et al. 2007; Zhuang et al. 2005) (**Figure 5E**). We crossed *Rorb*-Cre (Madisen et al. 2015), *Ntsr1*-Cre and *Slc6a4*-Cre lines to RCL-H2B-GFP reporter lines (Matho et al. 2021) and measured the spatial distributions of GFP expressing cells. Each line showed non-uniform areal and laminar specificity in cortex (**Figure 5F**). *Rorb*-Cre exhibited dense labeling in layer 4 of primary sensory cortex, but transitioned to labeling deeper layers in secondary sensory and primary motor cortex (Harris et al. 2014; Yamawaki et al. 2014)(**Figure 5G-I**). *Ntsr1*-Cre has been published as a layer 6-specific line, but labeled cells in dorsal insular cortex were in superficial layers, whereas prelimbic cortex showed a bimodal distribution of labeled cells (**Figure 5H, I**). *Slc6a4*-Cre labels cells that express *Slc6a4* transiently during development (Narboux-Nême et al. 2008). In cortex, we observed preferential labeling in midline and non-sensory cortices (**Figure 5H**). Labeled cells were biased toward deep layers in anterior cortex, but were bimodally distributed in posterior retrosplenial cortex and surrounding areas (**Figure 5I**). Brain-wide characterization of these transgenic mice reveals complex expression patterns that shift across the cortical sheet.

### Atlas-registered connectivity tracing with viral cytosolic labeling

We next demonstrate 3D-MAESTRO for mapping connectivity using cytosolic labeling of somata and neurites. We investigated the projection topography of locus coeruleus (LC). In *Dbh-*Cre:Ai65D mice, we performed retrograde injections of AAVrg-DIO-FlpO in either the main olfactory bulb (MOB) or between spinal cord vertebrae C4-C5 (**Figure 6A**). We used our cell detection model to localize the positions of labeled cells in 3D image volumes (Su et al. 2026). Projections to MOB were localized to superficial LC (**Figure 6B-E**). Projections to spinal cord occupied a larger volume of LC, but were biased ventrally (**Figure 6B-F**).

We used 3D-MAESTRO and cell segmentation to delineate hippocampal circuitry. We injected AAV1-mTurquoise into dentate gyrus (DG) and CA1 to label the input and output nodes of hippocampus (**Figure 6G, H**). AAV1 undergoes anterograde transcellular spread when delivered at high titer (Zingg et al. 2017; Faress et al. 2023). We observed labeled somata in DG, CA1, and subiculum (SUB), consistent with transcellular spread from CA1 to SUB (**Figure 6I, J**). To quantify the connectivity of these regions, we segmented neurites using a simple intensity threshold and then measured the masked volume in different CCFv3.1 regions (**Figure 6H-K**). We observed substantial labeling in CA3, as well as smaller labeled volumes in known targets of CA1 and SUB, including the ventral retrosplenial cortex (RSPv), postsubiculum (POST), rostral lateral septal nucleus (LSr), and deep layers of entorhinal cortices (ENTm and ENTl) (**Figure 6H, J, K**). Preferential transcellular spread from CA1 to SUB over other connected regions could reflect the strength of connectivity between these circuit nodes (Zingg et al. 2020).

**Figure 6.**
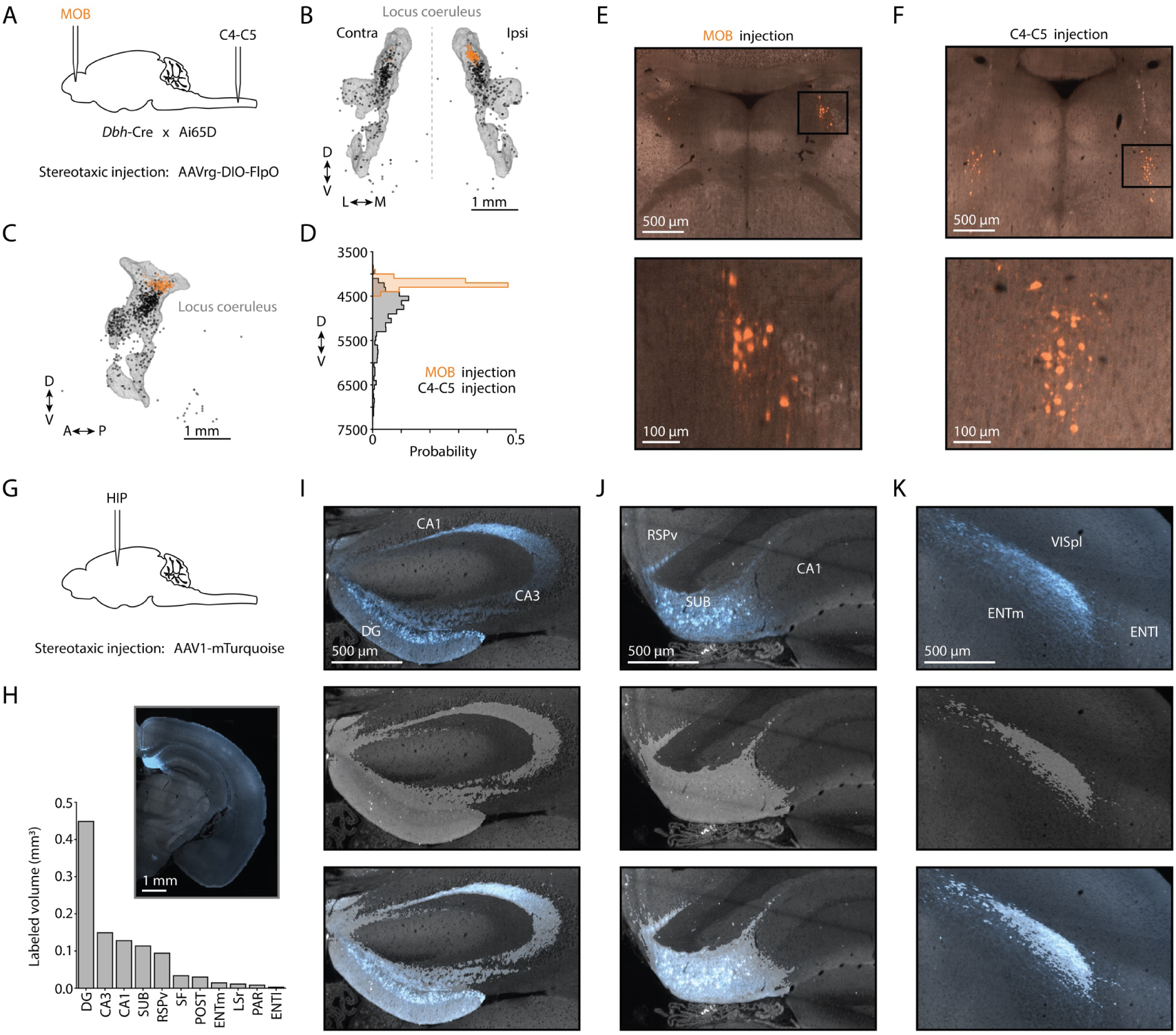
Atlas-registered connectivity tracing with viral cytosolic labeling. **A.** Stereotaxic injection of AAVrg-DIO-FlpO in either main olfactory bulb (MOB, *orange*) or spinal cord (C4-C5, *black*) of *Dbh*-Cre and RCL-tdTomato-WPRE (Ai65D) crossed animals. **B.** Coronal view of detected somata in CCFv3.1 space colored by injection, overlaid on a locus coeruleus mesh. **C.** Same as **B**, but sagittal view. **D.** Histogram of dorsal ventral positions of detected somata. **E.** Coronal view of locus coeruleus from MOB injected sample. **F.** Same as **E** for spinal cord injected sample. **G.** Stereotaxic injection of AAV1-mTurquoise in hippocampus. **H.** Segmented volume in CCFv3.1 regions. Inset, coronal view of injection site. **I.** Coronal view of hippocampus and dentate gyrus with fluorescent signal (*top*), segmented mask (*middle*), and merge (*bottom*). **J.** Same as **I**, but for subiculum. **K.** Same as **I**, but for entorhinal cortex. CA1 – field CA1, CA3 – field CA3, DG – dentate gyrus, ENTl – entorhinal area, lateral part, ENTm – entorhinal area, medial part, ENTm – entorhinal area, medial part, LSr – lateral septal nucleus, rostral part, PAR – parasubiculum, POST – postsubiculum, RSPv – retrosplenial area, ventral part, SF – septofimbrial nucleus, SUB – subiculum, VISpl – posterolateral visual area.

**Figure 7.**
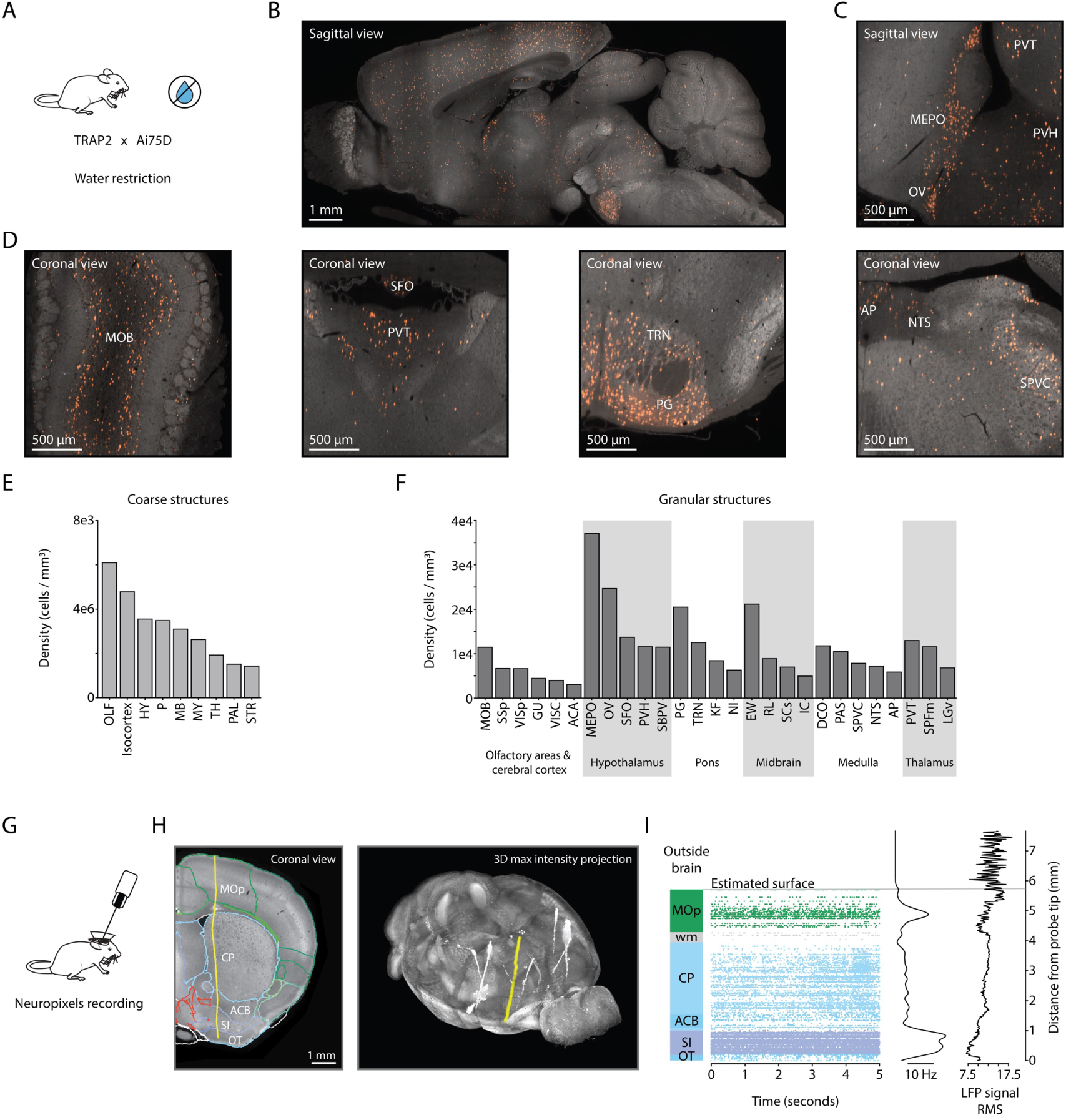
Neural activity maps. **A.** Targeted recombination in active populations (TRAP) in RCL-nls-tdTomato-WPRE (Ai75D) reporter mouse following water restriction. **B.** Sagittal view of whole mouse brain sample. **C.** Sagittal view of circumventricular nuclei. **D.** Coronal views of (*from lek to right*) main olfactory bulb, thalamus, ventral pons, and medulla. **E.** Detected cell density in coarse brain divisions. **F.** Detected cell density of granular brain structures. **G.** Experimental schematic of Neuropixel recording in the mouse brain. **H.** Coronal view of probe tract reconstruction in yellow (*lek*) and 3D max intensity projection of whole brain (*right*). **I.** Atlas-aligned spike recordings and LFP signal. ACA – anterior cingulate area, ACB – nucleus accumbens, AP – area postrema, CA – field CA, CP – caudoputamen, DCO – dorsal cochlear nucleus, EW – Edinger-Westphal nucleus, GU – gustatory areas, HY – hypothalamus, IC – inferior colliculus, KF – Kölliker-Fuse subnucleus, LGv – ventral part of the lateral geniculate complex, MB – midbrain, MEPO – median preoptic nucleus, MOB – main olfactory bulb, MOp – primary motor area, MY – medulla, NI – nucleus incertus, NTS – nucleus of the solitary tract, OLF – olfactory areas, OT – olfactory tubercle, OV – vascular organ of the lamina terminalis, P – pons, PAL – pallidum, PAS – parasolitary nucleus, PG – pontine gray, PVH – paraventricular hypothalamic nucleus, PVT – paraventricular nucleus of the thalamus, RL – rostral linear nucleus of the raphe, SBPV – subparaventricular zone, SCs – superior colliculus, superficial layer, SFO – subfornical organ, SI – substantia innominata, SPFm – subparafascicular area, medial part, SPVC – spinal nucleus of the trigeminal, caudal part, SSp – primary somatosensory area, STR – striatum, TH – thalamus, TRN – tegmental reticular nucleus, VISp – primary visual area, VISC – visceral area, wm – white matter.

### Neural activity maps

Linking measurements of neural activity to connectivity and gene expression requires registering multiple types of measurement to a common reference frame (Liu et al. 2021; Q. Wang et al. 2020).We used targeted recombination in active populations (TRAP) to perform a brain-wide screen for neurons expressing *Fos*, an activity-dependent immediate early gene, following water restriction (**Figure 7A**) (Luo et al. 2018; Ueta et al. 1995; Allen et al. 2017). We crossed *Fos*-CreERT2 mice and nucleus-localized tdTomato reporter mice. 4-Hydroxytamoxifen was administered after water restriction. 3D-MAESTRO was used to localize labeled cells (**Figure 7B-D**).

Subcortical regions showed the highest labeling density (**Figure 7F**), including known circumventricular thirst centers such as the vascular organ of the lamina terminalis (OV), subfornical organ (SFO), and area postrema (AP). Some of their efferent targets in the median preoptic nucleus (MEPO), paraventricular hypothalamus (PVH), and nucleus of the solitary tract (NTS) also showed prominent labeling (**Figure 7C, D**). Other subcortical areas linked to stress responses, such as the Edinger-Westphal nucleus (EW), paraventricular thalamus (PVT), nucleus incertus (NI) also showed activation.

Localizing electrophysiological recordings is critical for interpreting neural dynamics in the context of neural circuits. Neuropixels probes enable the recording of many neurons deep in the brain. Common practice is to coat probes in fluorescent dye prior to recording, then reconstruct probe tracks in histology (Laboratory et al. 2025; Liu et al. 2021). We demonstrated the use of 3D-MAESTRO for probe track reconstruction by estimating the location of units deep in the mouse brain (**Figure 7G**). Prior to recording, Neuropixels probes were dipped in either cm-DiI or sulfonate-DiD (Liu et al. 2021). We imaged the mouse brain and manually reconstructed probe tracks using Neuroglancer (**Figure 7H**). We used electrophysiology features (LFP spectra and correlation, unit locations, etc.) to estimate the location of specific electrodes along the probe track (**Figure7I**) (Laboratory et al. 2025). This mapping provides a precise measurement of the CCFv3.1 location for each recorded unit (Liu et al. 2021).

### Maps of inputs to brain regions based on retrograde viral labeling

We next evaluated the capacity of 3D-MAESTRO for measuring inter-areal connectivity in a scalable and reproducible manner. We performed stereotaxic injections of AAVrg (Tervo et al. 2016) expressing nucleus-localized fluorophores in cingulate or motor cortices (**Figure 8A**). To assess the variability in stereotaxic labeling we estimated the location and volume of each injection. Somata at the center of an injection site experienced a higher multiplicity of infection and expressed more fluorophore than cells retrogradely labeled via their axons. We modelled the fluorescence intensity at each injection site as a 3D Gaussian and estimated injection sites as a volume contained by the surface defined by 1/e of max intensity (**Figure 8B**). We used 3D-MAESTRO to obtain brain-wide maps of retrogradely labeled neurons. The patterns of labeling were qualitively different for cingulate and motor cortex injections (**Figure 8C**). The number of detected cells correlated with the estimated injection volume (**Figure 8D**). Hierarchical clustering of brain-wide input maps revealed that cingulate cortex and motor cortex injections clustered within groups, but differed greatly across groups (**Figure 8E**).

Both cortical areas received similar proportions of input from isocortex, but the specific input areas differed (**Figure 8F**). Cingulate cortex received greater input from association areas such as retrosplenial cortex and posterior parietal association areas, whereas motor cortex received greater input from somatosensory and interoceptive areas. Although thalamus provided a greater proprotion of input to cingulate compared to motor cortex, specific thalamic nuclei, such as the paracentral thalamus, provided more input to motor cortex. We consistently observed a small number of labeled cells in small neuromodulatory structures, including superior central raphe and ventral tegmental area, reflecting replicable CCFv3.1 registration between brains.

To assess the fine-scale spatial alignment of input maps, we computed 3D spatial histograms at 100 µm^3^ resolution from the locations of detected cells. We calculated the cosine similarity as a function of the distance between the CCFv3.1 location of the injection site (**Figure 8G,H**). More distant injections were less similar to one another. This relationship held both within and between injection areas, reflecting sampling along topographic gradients of cortical input. Thus we demonstrate 3D-MAESTRO’s capacity to enable scalable and reproducible investigation of neueral circuits at the level of the whole brain.

**Figure 8.**
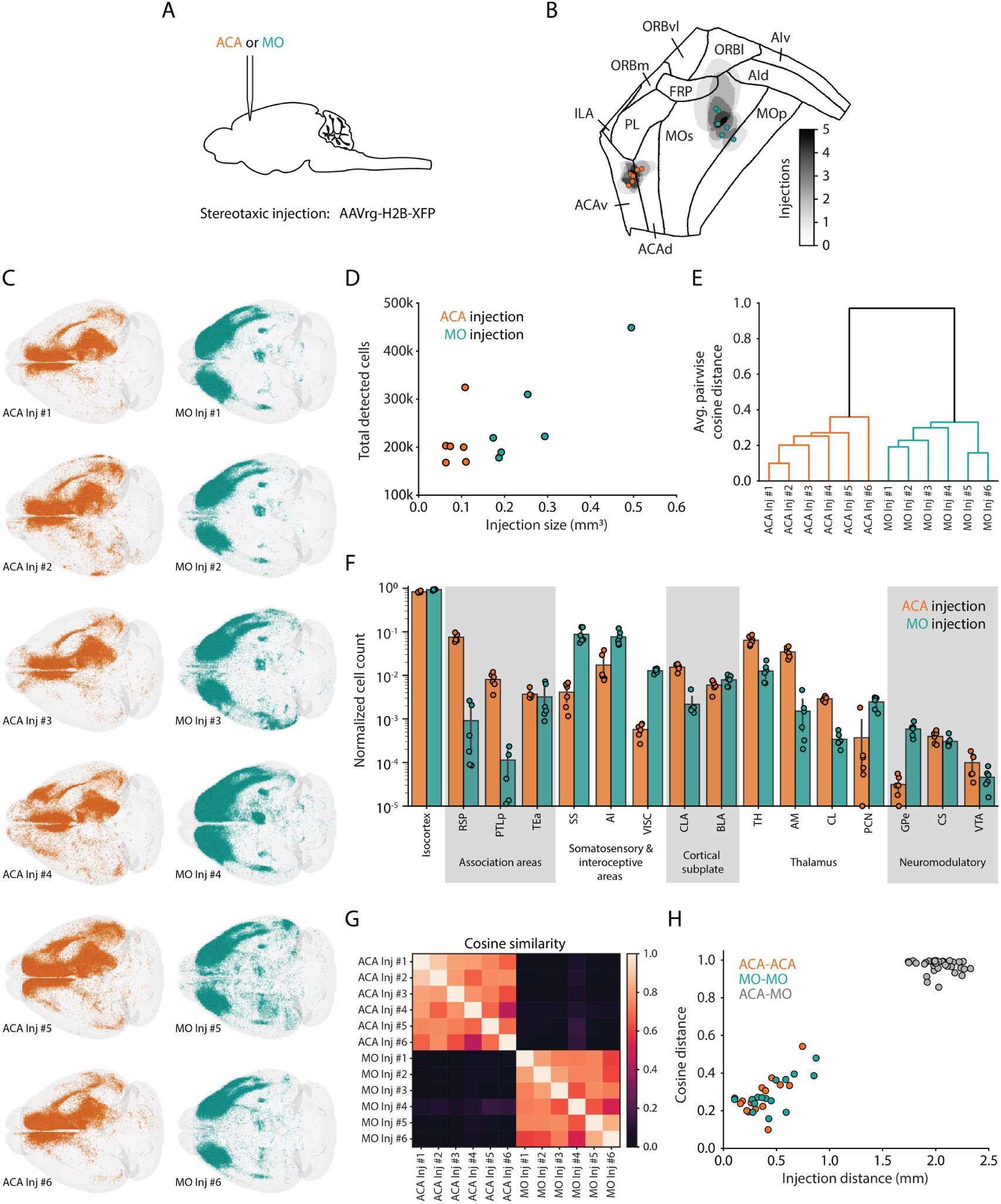
Maps of inputs to brain regions based on retrograde viral labeling. **A**. Stereotaxic injection of AAVrg-H2B-GFP or AAVrg-H2B-tdTomato in either cingulate cortex (*ACA, orange*) or motor cortex (MOs, *teal*). **B.** Representation of injection sites on a cortical flat map. Injection volumes were fit as 3d Gaussian profiles in CCFv3.1 coordinates. Centroids are marked by colored circles. Grey shading indicates the number of overlapping injections. **C.** Brain-wide maps of labeled neurons in retrograde labeling experiments. **D.** Total number of input cells detected as a function of injection volume. **E.** Hierarchical clustering of input maps. **F**. Region-wise input cell count, normalized to total cell count. **G.** Cosine similarity between brain-wide input labeling. 3D spatial histograms were computed in 100 µm^3^ voxels. **H**. Cosine distance of input maps as a function of distance between injection centroids. ACA – anterior cingulate cortex, MO – motor cortex, RSP – retrosplenial cortex, PTLp – posterior parietal cortex, TEa – temporal association cortex, SS – somatosensory cortex, AI – agranular insular cortex, VISC – visceral cortex, CLA – claustrum, BLA – basolateral amygdala, TH – thalamus, AM – anteromedial thalamus, CL – central lateral thalamus, PCN – paracentral thalamus, GPe – globus pallidus external segment, CS – superior central nucleus raphe, VTA – ventral tegmental area.

## Discussion

3D-MAESTRO is a robust, automated pipeline for the end-to-end processing of large-scale 3D microscopy data (**Figure 1**). The increasing scale, resolution, and molecular sophistication of large-scale 3D fluorescence microscopy promise a new era of *in situ* cellular and molecular biology. Capitalizing on these developments requires the ability to detect and measure small image features reliably across massive 3D volumes, and to do so in a consistent and reproducible manner in hundreds or thousands of specimens, across laboratories and image modalities. Experimental studies relying on dose–response series, panels of genetic reagents, cohorts spanning genotype, age, and behavioral condition, make automation a precondition for the science.

3D-MAESTRO manages the entire workflow from data ingest through quantitative, atlas-referenced output (**Figure 1**): it ingests standard image and metadata files, performs artifact correction, precise tile alignment and fusion and scalable analysis (e.g. atlas registration and cell counting). Intermediate products are inspectable in the browser via Neuroglancer. The pipeline is compatible with cloud infrastructure, on-premises HPC, and workstations. 3D-MAESTRO balances experimental adaptability against the requirement for rigorous reproducibility and data provenance.

Composing the pipeline in modular Nextflow workflows, with each stage packaged as a versioned container and each interface informed by a metadata schema, allows algorithms to be swapped and benchmarked in isolation (Table 4). For example, we used this feature to replace TeraStitcher with BigStitcher after head-to-head comparison (Section) and to test and deploy successive revisions of the cell-classification models. A pipeline that can absorb new algorithms in a modular fashion does not become obsolete as the field advances.

We applied 3D-MAESTRO to mapping the locations of mouse brain cells (**Figure 4**), registered to the CCFv3.1 mouse brain atlas (**Figure 3**). Mapping reporter gene expression based on enhancer AAVs and transgenic reporter mice is critical to evaluating these tools. Registering enhancer-AAV and transgenic reporter expression to the CCFv3.1 places genetic access tools in the same frame as the anatomical and transcriptomic taxonomies that define the cell types they are meant to target. Brain-wide maps of labeled cells reveal both on-target and off-target expression, providing critical information for using and improving these tools. Quantitative brain-wide maps often reveal departures from established descriptions. For example, Ntsr1-Cre labeling positioned superficially in dorsal insular cortex and bimodally in prelimbic cortex. These patterns are readily apparent when laminar depth is measured brain-wide in atlas coordinates rather than in selected sections. Retrograde and anterograde tracing yield connectivity in the same coordinates. TRAP labeling after water restriction places a functional variable in that frame. Reconstructed Neuropixels probe tracts anchor electrophysiological units, recorded in a behaving animal, to the same anatomical space as the molecular and connectional measurements. Each of these is informative alone; their value compounds when they are expressed in shared coordinates, because a structure can then be described simultaneously by what it is made of, what it connects to, and what it does. Building that composite picture is the principal scientific return on investment in registration accuracy.

Registration accuracy is an underappreciated limiting factor in whole-brain mapping. This is because registration algorithms degrade when contrast mechanisms differ between the sample and the reference. We demonstrate how building a modality-specific template can be used to achieve automated accurate registration of thousands of specimens to the CCFv3.1.

Overall, the value of 3D-MAESTRO lies less in any single component than in what the composition of these components makes routine. Over two years, 3D-MAESTRO has processed approximately 4,000 whole-brain datasets acquired on two SPIM microscopes, with turnaround of less than a day. Operation at this volume has itself been an engine of refinement: failure modes that appear rarely are invisible in a small study, but unavoidable at scale, and the pipeline is robust because it has been repeatedly repaired against real data.

Sustained throughput changes what can be asked. When whole-brain mapping is relatively routine, *n* becomes a design parameter rather than a constraint, and questions that were anecdotal become statistical. Our retrograde input-mapping experiments illustrate this shift: across twelve injections processed identically, we could relate labeled-cell counts to independently estimated injection volumes, cluster brain-wide input maps by target, and recover a topographic gradient in which input similarity declines smoothly with distance between injection centroids. Similarly, consistent detection of sparse labeling in tiny structures such as the locus coeruleus is enabled by aggregating experiments across animals, enabled by accurate registration.

3D-MAESTRO is designed to generalize beyond the mouse brain. The ingest, artifact-correction, alignment, and fusion stages are agnostic to tissue type and to microscope; the modality- and organ-specific commitments are confined to registration, segmentation, and detection, each of which is a customizable or replaceable module. The same infrastructure should therefore support whole-organ and whole-organism phenotyping, developmental time series, and comparative work across species. FAIR-aligned formats, machine-readable provenance for every processing step, and containerized environments are what allow datasets generated in different laboratories, on different instruments, and in different years to be pooled, compared and integrated. Fragmented, machine-specific workflows do not merely slow analysis; they silently foreclose the aggregation on which population-scale organismal biology depends.

Whole-brain, whole-organ, and eventually whole-organism imaging at cellular resolution is becoming a routine experiment. What determines whether it becomes a source of insight is whether the resulting data can be processed reproducibly, at volume, and placed in a frame where independent measurements can be compared. 3D-MAESTRO is our attempt to supply that infrastructure, and to do so in a form the community can build on.

Looking ahead, we aim to improve the performance of pipeline components, make them more configurable, and handle additional applications. For CCFv3.1 registration, we plan to add additional templates for widely used histology methods and imaging platforms. For cell detection, we are exploring the use of emerging 3D foundation models, which have the potential to substantially improve generality without loss of accuracy. We are developing a new Python codebase for large-scale, highperformance image stitching and fusion, which will support community contributions to leverage collective advancements in this area. Finally, to broaden accessibility and adoption across the neuroscience community, we are investigating the possibility of data processing services that would enable external researchers to use the pipeline in a seamless manner.

## Methods

### Data acquisition

Whole mouse brain samples were collected after cardiac perfusion and underwent, fixation, aqueous delipidation, index matching, and embedding as described in detail on protocols.io (https://www.protocols.io/workspaces/allen-institute-for-neural-dynamics/publications) (Allen Institute for Brain Science 2026; Myers & Toglia 2025; Myers & Toglia 2025; Myers & Toglia 2025). Imaging was performed using customized commercial SPIM microscopes (LifeCanvas SmartSPIM) with a 3.6×, 0.2 NA objective (Thorlabs/LifeCanvas) and voxel dimensions of 1.8 × 1.8 × 2.0 µm in xyz (Rohde 2023; Rohde 2023). Samples were imaged at 445, 488, or 561 nm excitation wavelengths, dependent on the fluorophores used in a given experiment. Background autofluoresence was imaged at 639 excitation wavelength.

Acquisition software was configured to disable automatic darkfield and flatfield corrections so that all image pre-processing could be performed within 3D-MAESTRO. Following data acquisition, custom scripts running on acquisition computers detected newly imaged datasets, collated and packaged the required metadata, verified data integrity, and transferred the datasets to an on-site storage system (VAST) with s3fs via s5cmd https://github.com/peak/s5cmd.

### Mouse surgery

All surgical and experimental procedures are similarly described on protocols.io (Stereotaxic injections (Allen Institute for Brain Science 2026), retro-orbital injections (Allen Institute for Brain Science 2026), electrophysiology surgery (Bennett et al. 2024; Lakunina et al. 2025; anna.lakunina 2024; Yin et al. 2024; Amaya et al. 2026), electrophysiology recording (Browning et al. 2026), water restriction (Amaya et al. 2024)). All procedures were in accordance with the National Institutes of Health Guide for the Care and Use of Laboratory Animals and approved by the Animal Care and Use Committees of the Allen Institute.

### SPIM template (ST)

This section outlines the development of the custom template for SPIM images of brains cleared in aqueous media, including manual structural annotations, and the mapping of the ST (**Figure 3G**) to the STPT (CCFv3.1 atlas; **Figure 3A,B**), which facilitate accurate sample registration. Here, we chose to construct a single-channel ST using autofluorescence from the far-red (639 nm excitation; 660-680 nm emission) channel, which is often a “free” channel without experimentally induced signal (3D).

Whole brain images tend to have uneven illumination intensities because of attenuation as the laser propagates through the tissue. The attenuation can differ across specimens, leading to inconsistent autofluorescence, which can cause registration errors. We therefore first performed intensity standardization across the entire stitched volume using the n4 algorithm (Tustison et al. 2010), with a large spline spacing of 1.5 mm. Next, we used Otsu thresholding (Otsu 1979) to create a mask. Applying the mask avoided out-of-brain artifacts from affecting registration or intensity normalization. All images were then normalized to the 98th percentile of signal in the image. This preprocessing is schematized in (**Figure S2A**). Because these intensity standardization steps were used in template construction, they must also be applied to any image being registered to the resulting template.

To construct the ST from the intensity standardized images, we employed a method similar to the ANTsPyX template building function. This function registers brains onto a common seed template, averages the results of these registrations, then updates the morphology of this average image by averaging the warping fields used to create it(2025; B. B. Avants et al. 2011). This process is performed iteratively, with each step serving as the seed template for the next step. We customized each step within this template construction framework to accommodate our large image volumes. Specifically, we moved each registration and morphology updating step to a designated worker and used NextFlow to parallelize the registration process across a pool of separate machines. We assumed the mouse brain to be symmetrical and registered each brain twice with one instance mirrored across its midline. Because occasional bright imaging artifacts can contaminate a final template, we used the median operation for the averaging step in place of the true average (mean) typically employed by ANTsPyX. **Table 2** details the genotype of the mice with samples selected for template construction. Each sample was manually inspected to ensure image quality in the 639 nm channel, and confirm that the brain was not damaged during histology.

**Table 2.** Mice used in ST construction.

| Mouse ID | Sex | Age (days) | Genotype |
| --- | --- | --- | --- |
| 679516 | M | 67 | Ai224(TICL-NLS-EGFP-ICF-NLS-dT)-hyg/wt |
| 685111 | F | 90 | Ai224(TICL-NLS-EGFP-ICF-NLS-dT)-hyg/wt |
| 693196 | F | 80 | wt/wt |
| 693197 | F | 66 | wt/wt |
| 693198 | M | 66 | wt/wt |
| 699091 | F | 91 | Dbh-Cre-KI/wt |
| 699396 | M | 88 | Dbh-Cre-KI/wt |
| 704365 | F | 59 | Nr5a1-Cre/wt;RCL-H2B-GFP/wt |
| 707207 | M | 59 | wt/wt |
| 713602 | M | 71 | wt/wt |

We initialized our pipeline using the median of all brains after a rigid registration to the STPT, ensuring that the result was in approximately the same anatomical space as this template. The “final” template was the output of five template-iterations. To preserve image contrast and detail, we constructed our template at 10 µm resolution. However, this created an unnecessary computation load for most individual sample registrations, and we therefore produced a 25 µm subsampling for STPT alignment.

In order to facilitate registration between our ST and the STPT, we selected the following structures for annotation small structures for annotation: IP/DN (the boundary between these areas was not clear, and so we grouped them together), LGd, RT, IPN, AD, act and III. Labels on the CCFv3.1 Atlas were obtained directly from the available annotations. To accelerate annotation of the ST, the CCFv3.1-space atlas labels were warped onto the ST using a mutual-information based direct registration with ANTs. Working from this starting point, our small structure annotations were hand curated ST, with the goal of producing structural drawings that matched as closely as possible the equivalent structure in the CCFv3.1 template atlas.

In addition we annotated the following large structures: CTX, CB, CNU, TH/MD/HB (again, when boundaries were not clear we grouped areas) and VS. For these, the CCFv3.1-space annotations were often modified to ensure correspondence between the two templates when details were ambiguous in the autofluorescence channel of either modality.

To compute the transform from ST to STPT template, we employed a hybrid registration approach that combined image-derived gradient information with manually annotated anatomical structures at both coarse and fine scales, defined in the coordinate spaces of each template (**Figure 3I**). Registration was performed using ANTs, enabling the joint incorporation of these complementary constraints. Mutual information (32 bins) was used to constrain image gradients, while mean squared error was applied to guide alignment of annotated structures. The gradient, small structures, and large structures received equal weighting in the deformable warp computation.

We evaluated our registration on individual brains by comparing manually drawn masks of the small structures listed above to the altas-registration output of our pipeline. Because the same structures were used to construct the template-to-template transform, we performed a leave-one-out procedure in which individual structures were excluded from the template-to-template transform computation so that they could be fairly evaluated. We quantified (1) the Dice score, that is, the percent overlap between to the warped CCFv3.1 mask for that structure and the hand-drawn annotation (**Figure S2 D**)(B. Avants et al. 2009) and (2) the average edge-to-edge distance between the outlines of both masks in 3D (**Figure S2 E**). Inter-rater Dice score in cases where the same structure was annotated by multiple people in the same mouse (n=4 structures) was 0.93, while the mean average surface-to-surface distance was 12µ*m*. Because our evaluation dataset was also used in constructing the template, this analysis was performed on brains included in the ST. Evaluation using a template that constructed from a separate set of wt/wt brains (albeit one built using fewer individual subjects) produced equivalent results (p<.001 for both metrics).

The SPIM template (ST), as well as an invertible mapping between the ST and the STPT (CCFv3.1), is available here: s3://aind-open-data/SmartSPIM-template_2024-05-16_11-26-14/.

### Cell detection

The network was trained on 59,721 annotations (23,399 true positives and 36,322 false positives), where true positives are points identified during proposal generation that were validated by an annotator to be a cell and false positive are point that were identified as not being a cell (e.g. tissue edge, nonspecific fluorescence, imaging artifacts). Annotations were collected across 30 mice (Male = 17; Female = 13; Age = 71 *±* 16 days) Seed locations for false positives were identified from proposals in the detection phase. The training data included images across various imaging channels, viral injections, and transgenic lines. Sample-wise percentile normalization (1, 99) and moderate augmentation was applied to compensate for the diversity of fluorescent signaling. Augmentations including changes in brightness, contrast, zoom, rotations, elastic deformation, and vertical/horizontal shifts, with each of the augmentations being applied independently with probability p = 0.1 per sample. This resulted in at least one transform being applied to roughly half of samples in a given epoch. This regiment was selected as we found it produced strong performance on our validation set. A cosine decay with reset schedule was used with a focal loss. Due to the imbalance in our training dataset, validation recall at precision 90 percent was monitored and the best weights across training epochs was saved. Model performance was evaluated on precision, recall and F1-score across nine benchmark datasets (Male = 3; Female = 4, Age = 67 *±* 10 days) with some individual animals having benchmarks across multiple channels.

Due to the variation in cell density and brightness a dynamic threshold for cell classification is calculated per dataset. During classification, the likelihood of each proposal being a cell is determined and a histogram of these values is created. The location of greatest separation between the two classed (i.e. global minimum) in the distribution is identified. Proposals with likelihood values above this point are classified as cells for quantification. This model is available here: s3://aind-benchmark-data/ tree/mesoscale-anatomy-cell-detection/models/smartspim_production_models/.

### Pipeline overview

The automated pipeline can be deployed on any system that is compatible with Nextflow. This includes the cloud (AWS), a local HPC or a local workstation. The pipeline starts with data ingestion. The ingestion step includes verification that the data adheres to pipeline standardization requirements. The pipeline then proceeds through six steps (**Figure 1**).

We next outline the data requirements and computational costs of the overall pipeline. Subsequent sections provide technical details related to each of the pipeline steps.

#### Pipeline input, output and parameters

The pipeline operates on multi-resolution multi-channel 3D microscopy datasets stored in OME-Zarr format. Required user-provided inputs are summarized in **Table 3**. The expected dataset structure is:

**Table 3.** Global inputs to the pipeline.

| Name | Type | Description |
| --- | --- | --- |
| DATA_PATH | Directory | Root dataset location containing raw OME-Zarr data and metadata |
| RESULTS_PATH | Directory | Destination for final processed outputs |
| template_path | Directory | Allen CCFv3.1 registration template |
| cell_models_path | Directory | Production deep-learning models for cell classification |

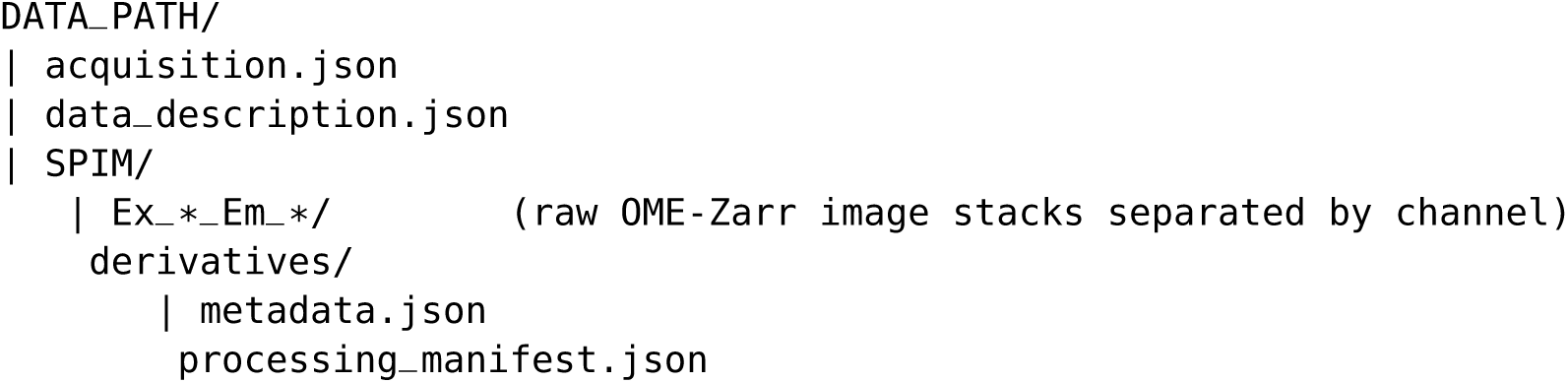

The pipeline produces the following user-facing outputs:

- Fused multiscale OME-Zarr image volumes
- CCFv3.1 registered images and spatial transforms
- Cell detection, classification, and spatial localization results
- Region-wise cell quantification tables
- Metadata and processing manifests (which outline the details of all the software processes and their versions applied to a dataset)

#### Computational cost

We estimated computational cost by mapping requested resources to equivalent on-demand cloud pricing (**Table 4**). These estimates contextualize resource requirements and are not exact billing values. Moreover, storage and data transfer costs depend on institutional infrastructure and dataset size and are therefore not included in the compute estimate.

**Table 4.** Resources, run time, and cost for all steps in the pipeline. Values are based on 6 datasets of sizes 875.07, 869.29, 882.78, 705.82, and 753.19 GBs. For steps marked as parallel (Y), multiple runs per dataset were executed (number of channels to preprocess and channels with cells to detect); the mean and standard deviation were first computed within each dataset, then combined across datasets to yield the reported values. Costs are calculated based on instance pricing. All values are reported as mean *±* standard deviation. *Cost per hour of 6 different whole-brain datasets with 3 channels each. Costs are estimated using the prices of the reported instance from https://aws.amazon.com/ec2/pricing/on-demand/ as of January 6, 2026.

| Step | Parallel? | CPUs | GPUs | RAM (GB) | VRAM (GB) | Run Time (h) | Instance | Cost* (USD) |
| --- | --- | --- | --- | --- | --- | --- | --- | --- |
| Vignetting correction (function estimation) | N | 4 | 0 | 32 | 0 | 4.57 ± 0.28 | r4.16xlarge | 1.22 ± 0.08 |
| Vignetting and stripe correction | Y | 16 | 0 | 128 | 0 | 5.46 ± 2.24 | r4.16xlarge | 22.13 ± 2.58 |
| Image Alignment | N | 16 | 0 | 256 | 0 | 1.33 ± 0.08 | m4.16xlarge | 3.48 ± 0.69 |
| Image Fusion | Y | 16 | 0 | 128 | 0 | 2.74 ± 0.61 | m4.16xlarge | 10.55 ± 1.60 |
| Atlas registration | N | 16 | 0 | 128 | 0 | 1.60 ± 0.23 | r4.16xlarge | 1.69 ± 0.24 |
| Job Dispatch | N | 2 | 0 | 16 | 0 | 3.37 ± 0.15 | r4.16xlarge | 0.45 ± 0.02 |
| Cell proposals | Y | 16 | 1 | 0 | 61 | 4.51 ± 0.74 | g4dn.4xlarge | 5.12 ± 0.87 |
| Cell classification | Y | 16 | 1 | 0 | 61 | 1.38 ± 0.55 | g4dn.4xlarge | 1.59 ± 0.66 |
| Cell quantification | Y | 16 | 0 | 128 | 0 | 0.86 ± 0.16 | r4.16xlarge | 0.99 ± 0.14 |
| Total | - | - | - | - | - | 25.82 ± 2.53 | | 47.22 ± 3.31 |

### Pipeline steps

This section outlines the implementation for the end user, assuming the availability of a reference SPIM template and a pre-trained cell classification model. While users may employ the assets provided in this work (e.g. the ST template and model for cell detection), the modularity of the pipeline enables these to be easily replaced.

Each module executes a discrete algorithmic task, generating standardized outputs, metadata, and images in OME-Zarr format to facilitate scalable whole-brain analysis. In addition, for downstream steps that can be run concurrently per channel (such as atlas registration and cell detection), we use the dispatch capability in NextFlow to reduce the wall time for data processing. Tasks that require all parallel runs to complete will wait for the aggregated results to start their execution.

The GitHub repository associated with this step is available at: https://github.com/AllenNeuralDynamics/aind-smartspim-pipeline-dispatcher

Because the framework is containerized and orchestrated via Nextflow, it is portable across diverse types of HPC and cloud environments. Resource requirements scale with data size to accommodate partial volumes, reduced channel counts, or other exploratory analyses.

#### Data Validation

Before data is processed, it is first validated:

1. Metadata files defined in the aind-data-schema package must exist:
  a. `acquisition.json`: defines microscope configuration parameters for data acquisition, including per-tile metadata such as resolution, orientation and nominal coordinates.
  b. `data_description.json`: defines administrative information such as the license, investigators, funding sources and institutions.
2. Folder structure (as described in the previous section : Pipeline Input, Output and Parameters)

The input for this step is the raw image data (OME Zarr format - using tools like ome-zarr-py, tensorstore, bioio) and metadata (aind-data-schema format). The output is a validation of whether the data formats are compliant with the pipeline. The GitHub repository associated with this step can be found at: https://github.com/AllenNeuralDynamics/aind-smartspim-validation

#### Image artifact correction

Once validated, the image data undergoes pre-processing (**Figure 1**) remove artifacts that compromise the robustness of downstream pipeline steps (**Figure 2**).

The input for this step are the acquired image tiles (OME Zarr format) and the output are image tiles with these artifacts removed (OME Zarr format). The GitHub repository associated with this step can be found at: https://github.com/AllenNeuralDynamics/aind-smartspim-destripe.

#### Stitching

These tiles then need to be “stitched” together to reconstruct a coherent volumetric representation of the sample before further analysis. Stitching is performed in two stages: tile alignment and fusion (**Figure 1**). During alignment, pairwise transformations are first estimated between overlapping images throughout the dataset. These local estimates are then integrated via a global optimization procedure to obtain a consistent alignment across all images (**Figure 2**). Image transformations are computed using BigStitcher (Hörl et al. 2019), which employs a phase-correlation algorithm at a resolution of 7.2 µm × 7.2 µm × 8.0 µm. The input for this step are acquired image tiles (OME Zarr format) and metadata (aind-data-schema format). The output is an alignment transformation XML file (Bigstitcher format). The GitHub repository for image stitching can be found at: https://github.com/AllenNeuralDynamics/aind-smartspim-stitch.

Using the computed alignment transforms, corrected tiles are fused into a multiscale volume in OME-Zarr format suitable for downstream analysis and visualization. We use BigStitcher and its OMEZarr writer, which generates a multiscale volume. The input for this step are image tiles (OME Zarr format), metadata (aind-data-schema format) and alignment transformation XML file (Bigstitcher format). The output is one fused volume (OME Zarr format). The GitHub repository for large-scale image fusion can be found at: https://github.com/AllenNeuralDynamics/aind-smartspim-fuse.

#### Atlas registration

The stitched image volumes are registered to the CCFv3.1 STPT and atlas via the ST. The channel that is selected for the alignment is specified by the user; channels close to the construction wavelength of the ST (639 nm) will work best because they are most similar to the ST itself, but alternative channels can be use if the 639 nm channel contains experimental signal that impairs registration (**Figure S2F**). The registration is estimated using this channel and then applied to all other channels.

The first step of the process is intensity standardization of the image volume (**Figure S3A**). This is needed to ensure intensity-scale consistency between the sample and the ST. Each sample is then registered into the ST space using ANTs with a neighborhood cross-correlation metric (radius 2). To account for differences in brain size and positioning, each sample underwent an initial rigid and affine registration before the fully deformable registration was run. Because the native image resolution of samples was higher than that of the template, registration was performed by resampling the Zarr level just below the 25 µm template resolution, typically 14.4µm x14.4µm x16µm. (ANTS inherently supports registration of objects with different specified resolutions.) We then applied the pre-computed template-to-STPT transform, yielding a complete registration from the input sample image to the Allen Common Coordinate Framework (CCFv3.1) atlas (**Figure 3J**).

The input for the Atlas Registration step was the fused volume (OME Zarr format), metadata (aind-data-schema) and CCFv3.1 (OME Zarr format). The outputs were a sample-to-CCF transformation files (ANTS format - .mat for the linear transforms and .nii for the warpfield). The GitHub repository associated with this step can be found at: https://github.com/AllenNeuralDynamics/aind-smartspim-ccf-registration.

#### Cell detection

The algorithm generates proposals of cell locations using a GPU-accelerated 3D Laplacian of Gaussian (LoG) algorithm. It accepts an OME-Zarr formatted volume and the algorithm is run block-wise at the highest resolution (level 0). The fluorescent profile of nuclei in SPIM images are generally elliptical (because of anisotropic resolution) with intensity peaks in the center. This makes them well suited for detection using conventional blob detection convolutions allowing for fast and accurate detection. (Matsumoto et al. 2019; Tyson et al. 2021; Toyoshima et al. 2016). Imaging volumes are lazily loaded and divided into super-chunks with volume size optimized to available GPU memory, minimizing transfer overhead. The super-chunk is then further divided into partially overlapping sub-volumes for detection (1A). During detection, background fluctuations are minimized across the volume to account for changes in auto-fluorescence as well as intensity transitions along tissue edges. This is done by calculating the 20th percentile of non-zero voxels and clipping the volume to the 20th percentile value. Next, a 3D LoG filter is applied to the processed image and peaks are identified through a combination of a maximum filter threshold on the filtered image and a raw threshold of the processed image. The LoG filter was optimized through a grid search over the parameter space. The combined threshold helps to eliminate peaks associated with noise in the image while maintaining a high level of sensitivity to low intensity signals. A common issue in conventional detection algorithms are the duplication of signals due to intensity fluctuations and clumping of densely packed cells (Toyoshima et al. 2016). To overcome these issues we pruned duplicate spots using a KD-Tree with a predefined context radius to remove duplicates and false detections along block edges. To further validate the identified spots a final gaussian fit is run on a volume crop centered around each spot’s centroid. A gaussian is fit across each axis and if convergence is not reached the spot is rejected (1B). The inputs for this step are the fused volume (OME Zarr). The output is `cell_likelihoods.csv` which contains the x, y, z location of all proposed cell locations, classification of proposal, likelihood, signal channel intensity, background channel intensity, and cell index. The GitHub repository associated with this step can be found at: https://github.com/AllenNeuralDynamics/aind-SmartSPIM-segmentation

After cell detection we apply a deep neural network binary classifier to separate true cells from potential imaging artifacts, such as auto-fluorescence or variation in regional tissue properties. We use a 3-D 18-layer ResNet backbone (He et al. 2015) applied to a 2x down sampled image for enhanced runtime performance. The network input is a (14, 14, 26, 2) block centered on a location from the detection phase containing both the signal channel as well as the background channel with only native auto-fluorescence (1C). The benefits of the detection-classification setup are two-fold. As mentioned above, this allows for removal of broad spectrum auto-fluorescence indicative of artifacts from tissue processing often missed in algorithms that do not take cross-channel signaling into account. Second, it allows for liberal detection parameterization minimizing the potential of false negatives. The inputs for this step are `cell_likelihoods.csv` and a pretrained classification model (keras format). The outputs are `detected_cells.csv` which contains the x, y, z location of only proposals that were classified as cells; and `cell_likelihood_metrics.csv` which contains high-level quantification of classification performance including the number of cells, mean likelihood, and likelihood standard deviation for both proposals that were classified as cells and not cells, and the dynamically set threshold; a thresh-old_identification.png showing a histogram of the proposal likelihoods, a smoothed curve fit to the data and the location of the threshold; annotations in neuroglancer’s precomputed annotation format and a `neuroglancer_configs.json` file for visualization classified cell locations overlaid on the fused image volume. The GitHub repository associated with this step can be found at: https://github.com/AllenNeuralDynamics/aind-smartspim-classification

#### Cell quantification

To quantify whole-brain and region-specific cell counts, detected cell coordinates are transformed from physical space into CCFv3.1. The transformation is implemented using the ANTsPy biomedical image processing toolkit (Tustison et al. 2021). Region-specific cell counts are derived using precomputed anatomical meshes covering 841 brain regions. Processing is parallelized across regions, yielding outputs that include total cell counts, regional volumes, and cell densities, calculated both within and across hemispheres. To assess the quality of registration, additional measurements of signal intensity and regional volumes are computed for a subset of eight brain areas. These metrics serve as confidence indicators and can help identify discrepancies or artifacts in the registration of both SPIM volumes and cell positions to the CCFv3.1.

The inputs to this step are detected_cells.csv which contains the locations of the detected cells and the transformations that map the sample into CCFv3.1 Space computed in Atlas registration (ANTs format: .mat file and .nii file). The output file (`cell_count_by_region.csv`) contains the number of cells in each CCFv3.1 region:

- `ID`: Unique numerical identifier for a given region
- `Acronym`: Acronym associated with region
- `Name`: Full name of region
- `Parent Region`: Location of region within commonly used high-level parent regions
- `Layer`: Cortical layer of region if applicable
- `Ancestors`: Full list of upstream parent regions
- `Graph Order`: Location within the Allen Brain Atlas structure graph
- `Struct_Info`: Identifies regions that across the midline or are separated in each hemisphere
- `Struct_area_um3`: The total volume of a region in voxels
- `Left`: Total number of cells identified in the left hemisphere of a region
- `Right`: Total number of cells identified in the right hemisphere of a region
- `Total`: Total number of cells identified within a region
- `Left_Density`: Density of cells (cells/mm^3^) identified in the left hemisphere of a region
- `Right_Density`: Density of cells (cells/mm^3^) identified in the right hemisphere of a region
- `Total_Density`: Total density of cells (cells/mm^3^) identified within a region
- `Left_Median_Foreground`: The median signal strength of cells identified in the left hemisphere of a region
- `Right_Median_Foreground`: The median signal strength of cells identified in the right hemisphere of a region
- `Total_Median_Foreground`: The median signal strength of cells identified in a region
- `Left_Median_Background`: The median background signal surrounding cells identified in the left hemisphere of a region
- `Right_Median_Background`: The median background signal surrounding cells identified in the right hemisphere of a region
- `Total_Median_Background`: The median background signal surrounding cells identified in a region

The output (transformed_cells.csv) contains cell coordinates in CCFv3.1 space:

- `x`: location in the Anterior-Posterior axis
- `y`: location in the Dorsal-Ventral axis
- `z`: location in the Medial-Lateral axis

The GitHub repository associated with this step can be found at: https://github.com/AllenNeuralDynamics/aind-smartspim-quantification

### Evaluation

#### Image artifact removal

To evaluate the correction for vignetting artifacts, we adopted two metrics. The first one is the intensity profile, which measures intensity changes along one dimension between the uncorrected and corrected images. The second metric is the Pixel Value Spread (PVS). To be able to properly evaluate the correction, we perform image thresholding creating a region of interest (ROI), where the metric is computed. We show axial slices before and after vignetting correction (**Figure 2a**). The quantitative metrics support the visual improvements: the intensity profile along the *y* axis becomes more uniform with the removal of the dips corresponding to the dark bands.

The Pixel Value Spread (PVS) is defined as follows:

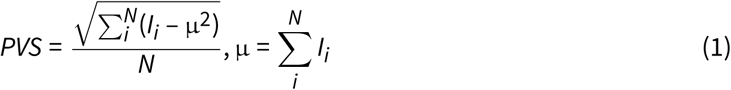

where index *i* iterates over pixels of an image *I*, *N* is the total number of pixels, *I_i_* represents individual pixel intensities, and µ is the mean pixel intensity. The PVS metric quantifies the standard deviation of pixel intensities in flat regions of the image. Lower PVS values indicate better correction quality, as reduced pixel-to-pixel variation in uniform areas corresponds to more successful removal of stripe artifacts and noise. Higher PVS values suggest greater intensity fluctuations and indicate the presence of uncorrected striping or noise artifacts.

In order to evaluate the presence of these stripes in images, we use the line difference metric (Varga 2020) which is defined as:

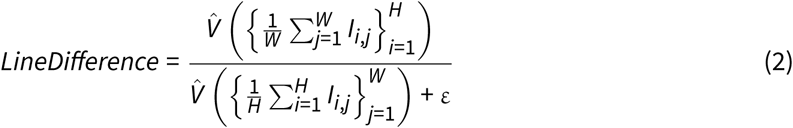

where *H* is the image height, *W* is the image width, *I_i,j_* is the pixel intensity at position (*i*, *j*), the numerator computes the variance (*V̂*) of row means, the denominator computes the variance of column means, and ε is a small positive constant to prevent division by zero. For horizontal stripe artifacts, each row has approximately uniform intensity, but different rows have significantly different mean intensities resulting in a high variance of row means (horizontal intensity variance). Considering that the other axis is not affected by the stripes (due to the orientation of the light sheet), there will be a low variance of the column means (vertical intensity variance). Therefore, values closer to 0 indicate fewer striping artifacts, and higher values indicate more artifact severity in the corresponding axis.

For evaluating the destriping algorithm, we extracted a block from a view oriented orthogonally to the dominant striping axis to use as ground truth that is largely free of striping artifacts. We then extracted stripe patterns from data exhibiting striping artifacts and artificially inserted them into this block. The resulting image was subsequently destriped and we computed the line difference metric for all three cases: 0.63 for the original stripe-free block, 0.70 for the block with artificially introduced stripes, and 0.63 for the corrected block. After destriping, the metric closely approaches the value of the original data, indicating effective removal of the introduced striping artifacts (**Figure 2b**).

#### Image tile alignment

To quantify alignment quality, we developed a cell-based error metric defined on pairs of overlapping image tiles. The metric compares two quantities: (i) the pairwise translation between each tile pair, measured directly from image content, and (ii) the per-tile transforms produced by global optimization and used for fusion. Each pairwise translation is estimated independently of every other pair and of the fitted transforms, so the discrepancy between the two measures how faithfully the fused volume reproduces the alignment implied by the image content, evaluated at the positions of detected cells. For tile *I*, denote the pre-alignment placement on the acquisition grid as *T_I_*, the fitted correction from global optimization as *A_I_*, and the full transform applied during fusion as *M_I_* = *A_I_T_I_*. (Here, affine transforms act on homogeneous coordinates.)

Measured pairwise shift: For each pair of overlapping tiles (*I*, *J*), the translation *s_IJ_* that best aligns them was estimated by phase correlation over the shared volume, together with a correlation coefficient quantifying confidence in that estimate. Shifts are defined in the pre-alignment frame, before any correction is applied, and are computed independently for every tile pair; they are therefore observations of the image data rather than products of the alignment model. The set of shifts forms the input to global optimization and is recorded in the alignment metadata.

Cell locations in image tiles: Cell detections obtained in the fused volume were assigned to tiles by inverting each tile’s full transform: cell *p* belongs to tile *I* when *M_I_*^−1^*p* falls within that tile’s voxel bounds. Cells assigned to two or more tiles form the validation set. Assignments were verified by manual inspection in Neuroglancer, confirming that positions back-projected into each tile coincided with the same cell in both tiles of the pair.

Inter-Image Cell Correspondence Error: Consider a cell at global position *p* that lies in both tile *I* and tile *J*. We locate it in each tile, then bring both back to the fused volume and measure how far apart they land. Undoing tile *I*’s correction gives its position as tile *I* recorded it, *q_I_* = *A_I_*^−1^*p*. The measured shift gives the same cell’s position as recorded by tile *J*, *q_J_* = *q_I_* – *s_IJ_*. Re-applying each tile’s correction maps both back into the fused volume, and the ICCE is the Euclidean distance between them in metric units:

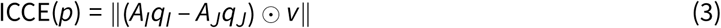

where voxel size in metric units is *v* = (*v_x_*, *v_y_*, *v_z_*) and ⊙ denotes element-wise multiplication. The measured shift pairs the cell across the two tiles, whereas the mapping into the fused volume comes from the fitted corrections; ICCE is therefore zero when the corrections reproduce the shift and grows as they disagree. Because the error is evaluated at cell locations rather than over the overlap as a whole, it is weighted by where the objects of interest actually lie, providing a biologically-relevant measure.

#### Cell detection and classification

For calculation of precision, recall and F1 scores, we first identified true positives (predictions with corresponding annotations), false positives (predictions with no corresponding annotations) and false negatives (annotations with no corresponding predictions). As our detection algorithm represents each identified object as a single coordinate point (centroid), there are many instances in which a cell can be correctly identified but not at the exact voxel location of the benchmarks annotation. To account for this, we use a distance-based threshold set a 14µm (7 voxels) between an annotation and prediction. This accounts for slight variations in where the algorithm placed the cells centroid relative to the expert annotator, while minimizing the likelihood of misidentifying a prediction from other cells. For each annotation, if the model prediction was within this distance it was assigned as a true positive. After a cell location from the benchmark was paired the annotation as well as the predicted point were removed from further analysis to avoid duplicate pairings. Additionally, if more than one prediction was within this threshold (duplicate labeling), the nearest point was paired to the annotation and removed, but the additional point was left. If no benchmark location was within 14µm, or if all locations within that range were already paired, a model prediction was labeled as a false positive. Conversely, any benchmark annotation that was remaining after all detected cells were paired, i.e., did not have an annotation within 14µm or annotations within range were already paired, were labeled as false negatives. For comparison with segmentation based algorithms, the centroid from individual segmentation masks were used to define the cell’s location and the above process was performed.

### Visualization

Image processing pipelines are commonly developed and executed on-premises, allowing researchers to visualize multi-dimensional biological datasets using powerful desktop applications such as ImageJ/Fiji (Schindelin et al. 2012), Napari (Ahlers et al. 2023), QuPath (Bankhead et al. 2017), and 3D Slicer (Pieper et al. 2004). These tools often lack native support for reading large-scale image data directly from cloud storage systems such as Amazon S3 or Google Cloud Storage. Several cloud-based visualization tools—such as Neuroglancer (2025), Horta Cloud (Rokicki et al. 2025), and OMERO Viewer—have emerged to address this limitation. In evaluating these tools, key criteria included efficient rendering of large OME-Zarr–formatted images, support for data located in cloud storage, support for visualizing spatial annotations (e.g., point clouds and surface meshes), and ease of sharing visualizations via URLs. We use Neuroglancer (2025), a browser-based tool developed to support large-scale electron microscope images, due to its performance, flexibility, and sharing capabilities. These are essential for enabling collaborative, cloud-native imaging workflows.

To facilitate integration with Neuroglancer, we automate the generation of shareable hyperlinks during pipeline execution. This supports multi-channel image visualization and the generation of Neuroglancer-compatible formats for spatial annotations, including point sets and meshes.

## Data and code availability

The pipeline and associated tools are available at: https://github.com/AllenNeuralDynamics/aind-smartspim-pipeline and https://github.com/AllenNeuralDynamics/aind-mesoscale-tools They all have been tested on standard workstations with a Linux platform and to enable the adaptation of this protocol to other types of imaging experiments, we provide the source code and documentation for each individual module as well as the raw and fused volume datasets that were processed with this protocol, which are available at https://s3.console.aws.amazon.com/s3/buckets/aind-open-data?region=us-west-2.

## Acknowledgments

This research was supported by the Allen Institute, founded by Jody Allen – chair and co-founder of Allen Family Philanthropies, and the late Paul G. Allen – investor, philanthropist, and co-founder of Microsoft. We gratefully acknowledge their vision and generosity, which make this work possible. Research reported in this publication was supported by the National Institute of Mental Health (NIMH) under award numbers UF1MH128339, U01MH139778 and RF1MH128841, the National Institute on Drug Abuse (NIDA) and National Institute of Neurological Disorders and Stroke (NINDS) under award number U19NS123714 and the AWS Open Data program. We thank Jada Roth, Emma Thomas, Tamara Zeric, Peter Grotz, Brooke Wynalda, Akira Fushiki, Rana Kutsal, Adrien Stanley, Kanghoon Jung, Jonathan Wong, and Zoe Juneau for helping us with data annotation for cell detection, as well as Hanna Belski, Corbette Bennett, Hannah Cabasco, Kevin Cao, Mikayla Carlson, Severine Durand, Akira Fushiki, Ryan Gillis, Han Hou, Kanghoon Jung, Gabor Kovacs, Ethan McBride, Arjun Sridhar, Adrien Stanley, and Xinxin Yin for helping annotate the validate dataset for the ST-to-STPT transform. We thank members of the Allen Institute (Neural Dynamics and Brain Science) for using the pipeline to process their datasets, and providing feedback on pipeline outputs and visualizations.

## Author contribution matrix

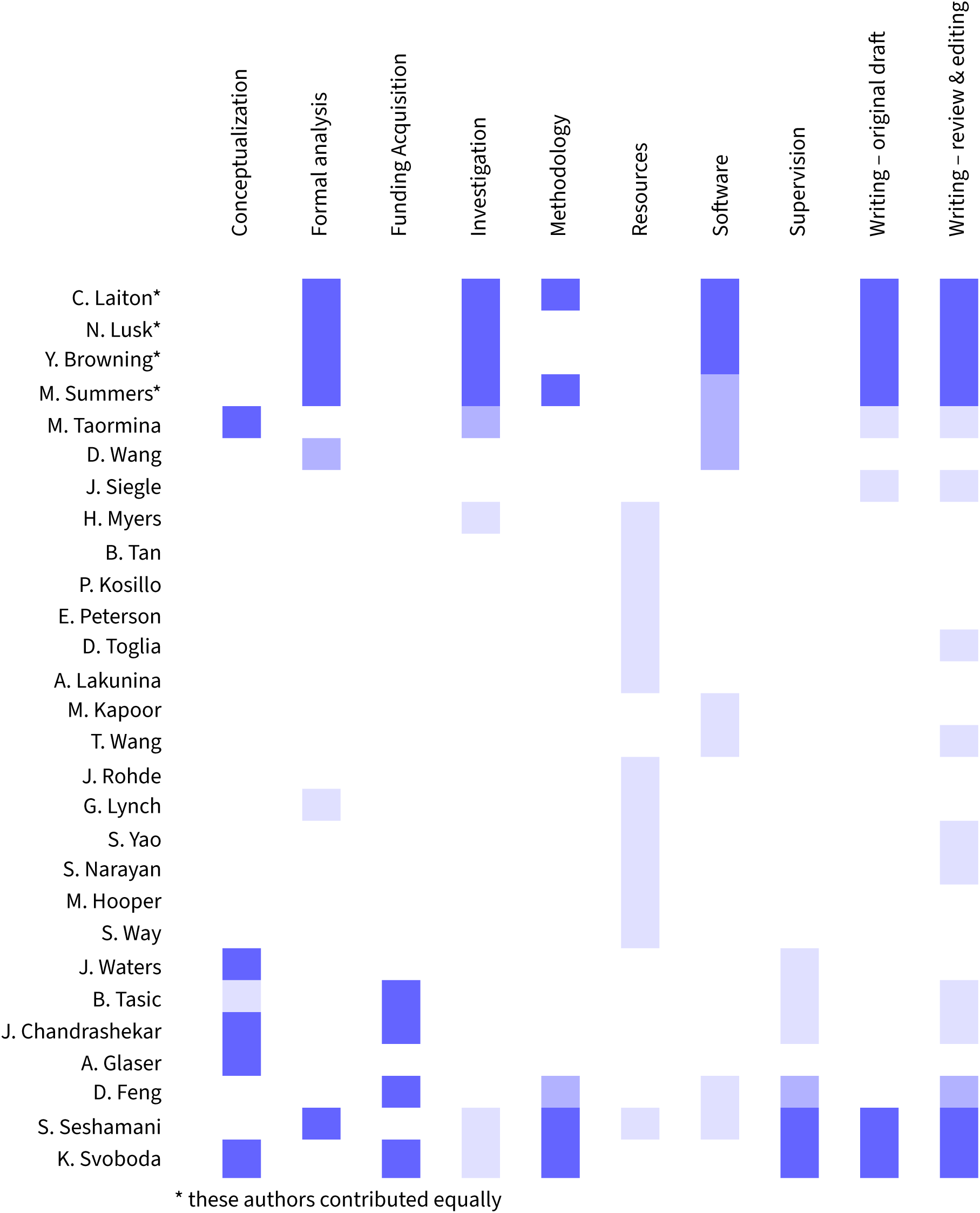

---

**Figure S1.**
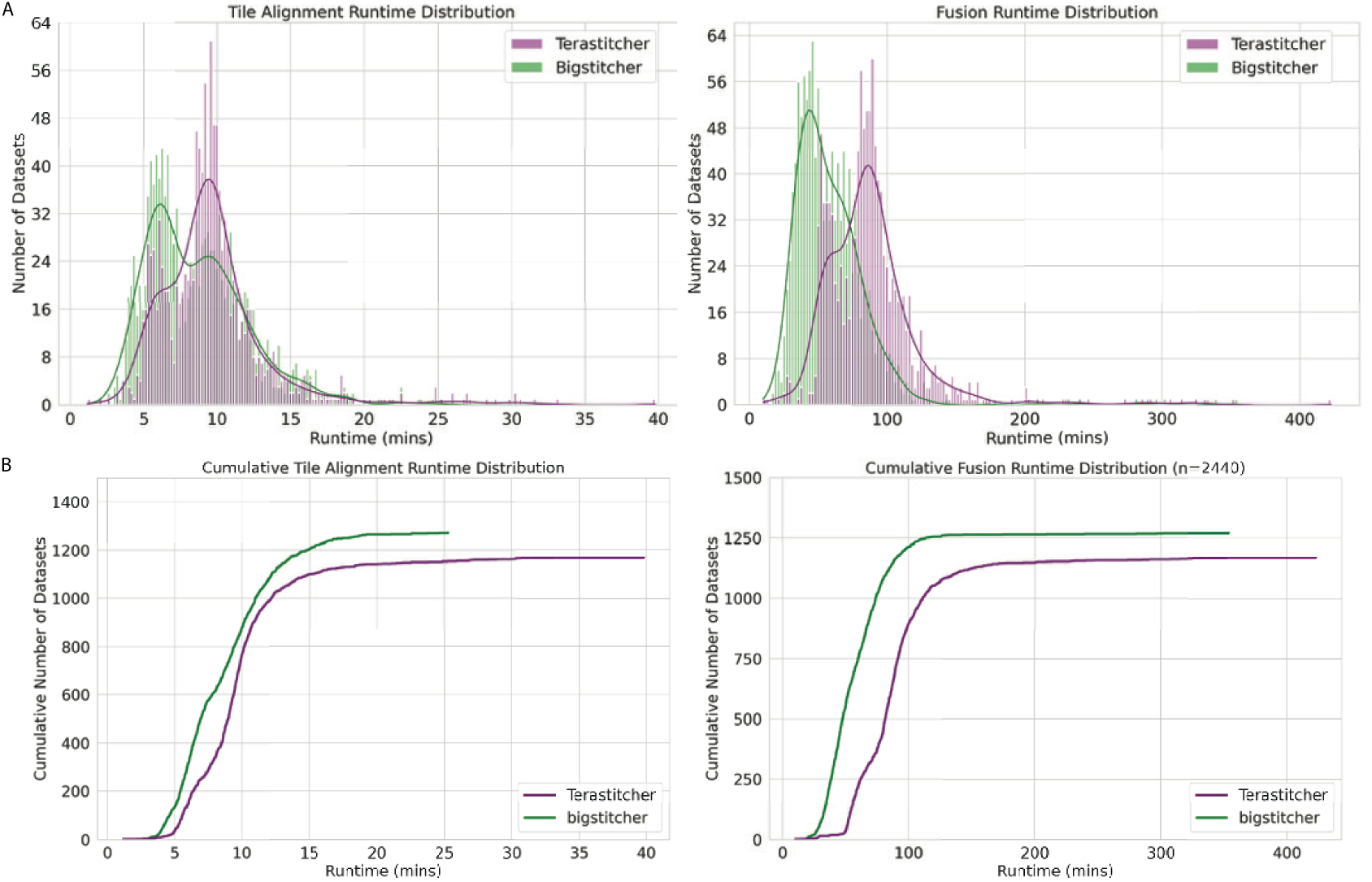
Stitching Comparison: **A,** Histogram of processing times for alignment and fusion for Terastitcher and Bigstitcher **B,** Cumulative Histogram of processing times for alignment and fusion for Terastitcher and Bigstitcher. Note that the maximum time for running Bigstitcher for Tile Alignment and Fusion is significantly lower for Bigstitcher than for Terastitcher, especially for fusion.

**Figure S2.**
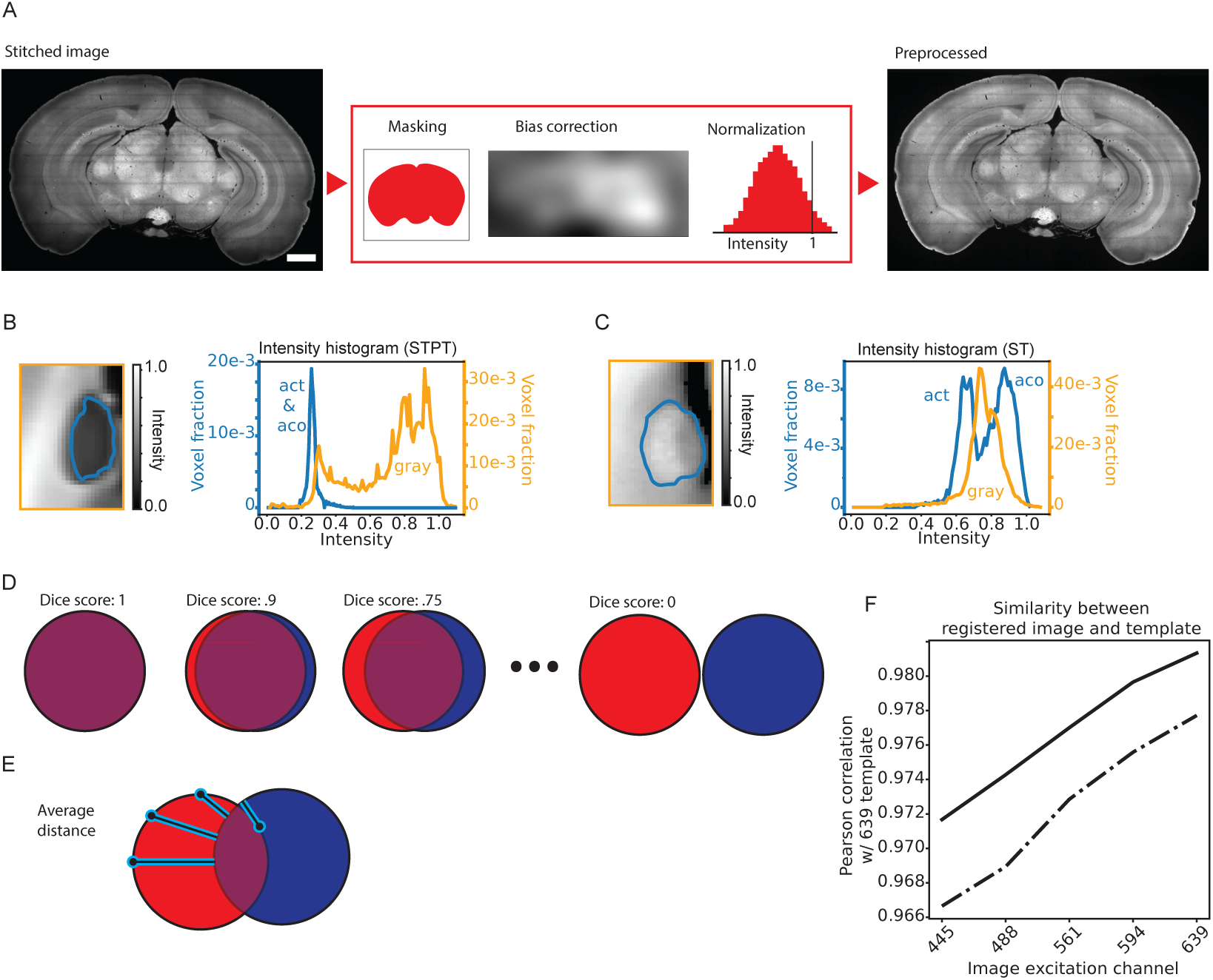
Atlas Registration. **A,** Preprocessing steps. A coronal slice from single specimen (639 nm) is shown before and after this preprocesing. **B,** (left) Cutout highlight the anterior commissure in the STPT from Figure 3C and (left) the image intensity within the highlighted areas. **C,** Same as *B,* but for the ST **D** Schematic illustrating Dice scores between the red and blue circles. **E** Same as *D*, but illustrating average edge-to-edge distance. **F** Pearson correlation between an individual brain and the an average brain. Each line indicates a different brain, registered using the 639 nm template but using different image channels.

**Figure S3.**
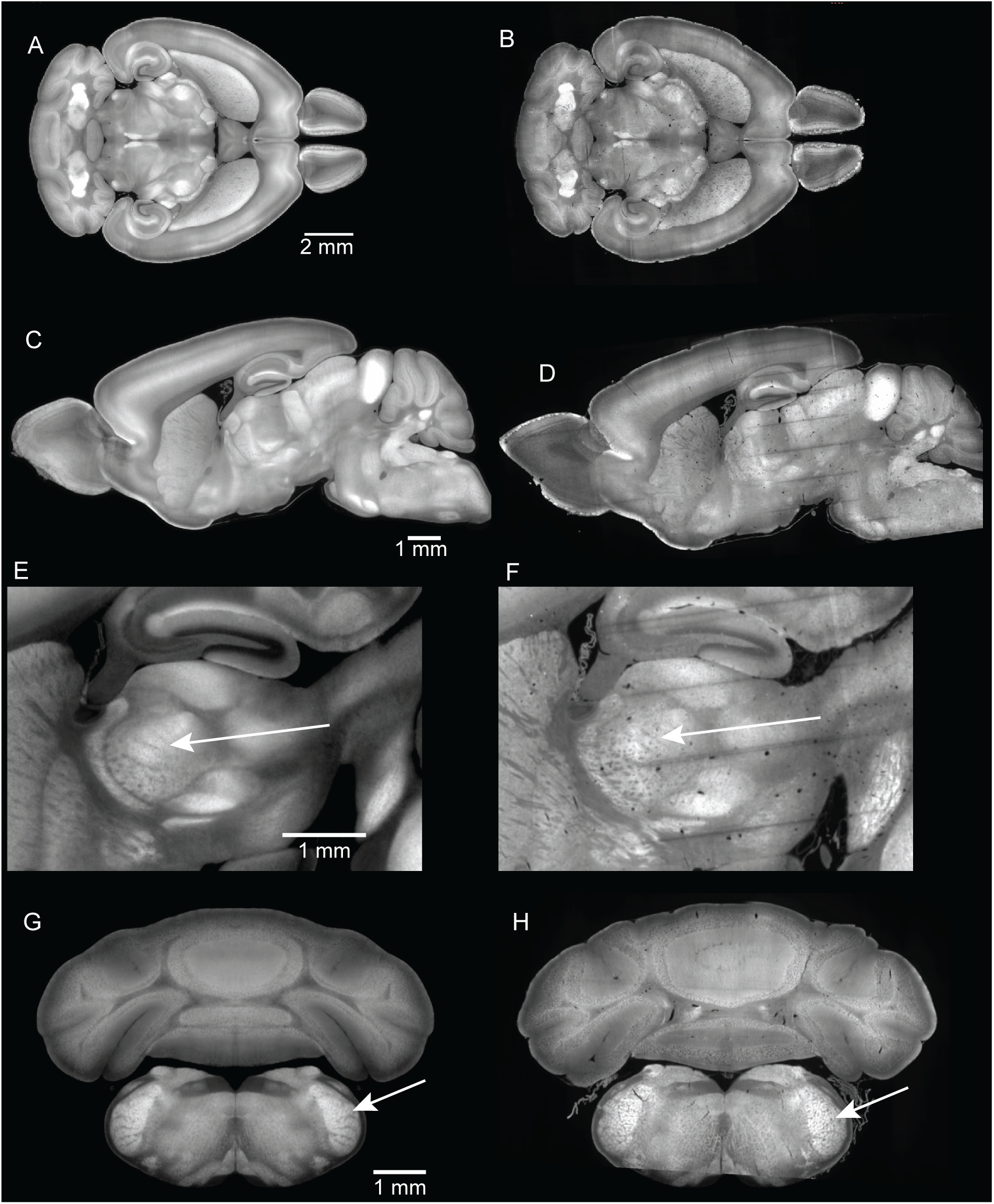
SPIM Template (ST). **A,** Temporal slice from the ST and **B,** Equivalent slice from a single specimen. **C,D,** Sagittal slices showing the whole brain **E,F,** Sagittal slices highlighting barreloids in ventral posteromedial nucleus of thalamus (VPM) G,H Sagittal slices highlighting barrelettes in the interpolar part of the spinal nucleus of the trigeminal nerve (SPVI).

**Table S1.**
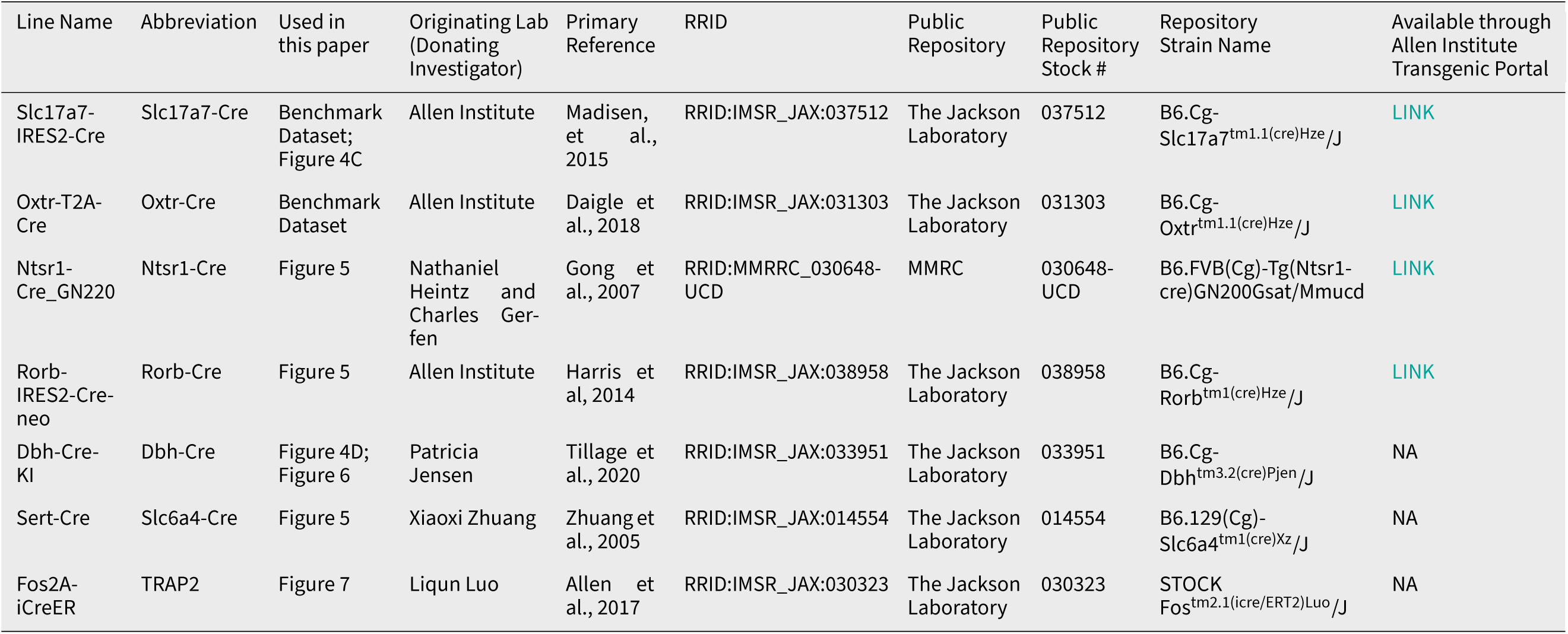
Driver Lines used in this study.

**Table S2.** Reporter Lines used in this study.

| Line Name | Abbreviation | Used in this paper | Originating Lab (Donating Investigator) | Primary Reference | RRID | Public Repository | Public Repository Stock # | Repository Strain Name | Available through Allen Institute Transgenic Portal |
| --- | --- | --- | --- | --- | --- | --- | --- | --- | --- |
| RCL-H2B-EGFP | RCL-H2B-EGFP | Benchmark Dataset; Figure 4C; Figure 5 | Z. Josh Huang | Matho et al., 2021 | RRID:IMSR_JAX:036761 | The Jackson Laboratory | 036761 | B6.Cg-Gt(ROSA)26Sor<sup>tm8</sup>(CAG-HIST1H2B | NA |
| RCFL-tdTomato-WPRE-D | Ai65D | Figure 6 | Allen Institute | Madisen, et al., 2015 | RRID:IMSR_JAX:021875 | The Jackson Laboratory | 021875 | B6.Cg-Gt(ROSA)26Sor<sup>tm75.1</sup>(CAG-tdTomato | NA |
| RCL-nls-tdTomato-WPRE-D | Ai75D | Figure 7 | Allen Institute | Daigle et al., 2018 | RRID:IMSR_JAX:025106 | The Jackson Laboratory | 025106 | B6.Cg-Gt(ROSA)26Sor<sup>tm75.1</sup>(CAG-tdTomato | NA |

**Table S3.** Viruses and viral constructs used in this study.

| Full Plasmid Description | Enhancer ID | Abbreviation | Used in this paper | Originating Lab (Donating Investigator) | Primary Reference | Plasmid ID | Public Repository | Public Repository Stock # |
| --- | --- | --- | --- | --- | --- | --- | --- | --- |
| pAAV-AiE0452h-minBG-SYFP2-P2A-mScarlet-10aa-H2B-WPRE3-BGHpA | AiE0452h | AiE0452h-SYFP2/mScarlet-H2B | Benchmark Dataset | Allen Institute | Hunker et al., 2025 | AiP15143 | Addgene | 224193 |
| pAAV-AiE2333m-minBG-SYFP2-P2A-mScarlet-10aa-H2B-WPRE-BGHpA | AiE2333m | AiE2333m-SYFP2/mScarlet-H2B | Figure 4A | Allen Institute | This Paper | AiP20274 | Addgene, in progress | Addgene, in progress |
| pAAV-AiE2252m-minBG-iCre(R297T)-BGHpA | AiE2252m | AiE2252m-iCre(R297T) | Figure 5 | Allen Institute | This Paper | AiP1837 | Addgene | 241591 |
| EnvA CSV-N2C-histone-GFP | NA | EnvA-H2B-GFP | Benchmark Dataset | UNC Vector Core (Custom Prep) | Yao et al., 2023 | NA | Na | Na |
| AAVrg-Syn-H2B-EGFP | NA | AAVrg-H2B-EGFP | Benchmark Dataset | UNC Vector Core (Custom Prep) | Tervo et al., 2016 (for capsid) | NA | Na | NA |
| AAVrg-Syn-H2B-Turquoise | NA | AAVrg-H2B-Turquoise | Benchmark Dataset | UNC Vector Core (Custom Prep) | Tervo et al., 2016 (for capsid) | NA | Na | NA |
| AAVrg-Syn-H2B-tdTomato | NA | AAVrg-H2B-tdTomato | Benchmark Dataset | UNC Vector Core (Custom Prep) | Tervo et al., 2016 (for capsid) | NA | Na | NA |
| AAVretro-DIO-FlpO | NA | AAVretro-DIO-FlpO | Figure 4D; Figure 6A-F | Li Zhang | Zingg et al., 2017 | NA | Addgene | 87306 |
| AAV1-mTurquoise | NA | AAV1-mTurquoise | Figure 6G-K | UNC Vector Core (Custom Prep) | Zingg et al., 2017 (for capsid) | NA | NA | NA |

**Table S4.**
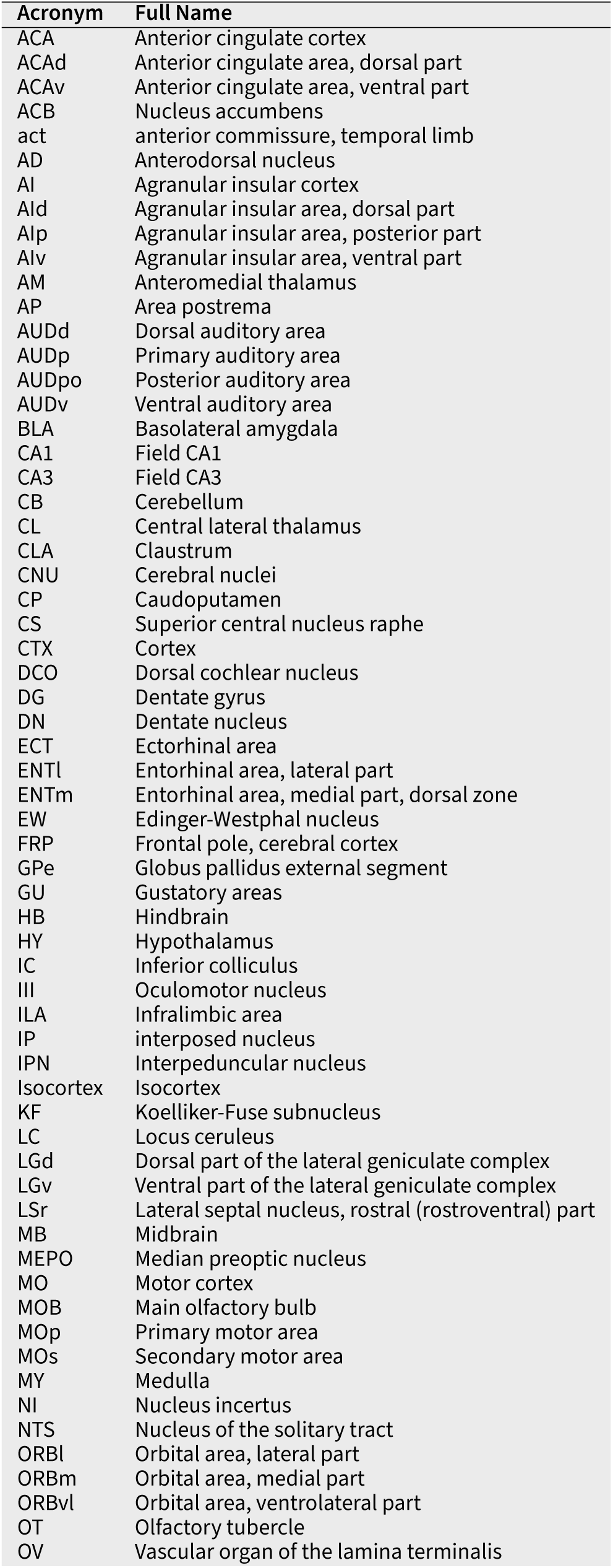
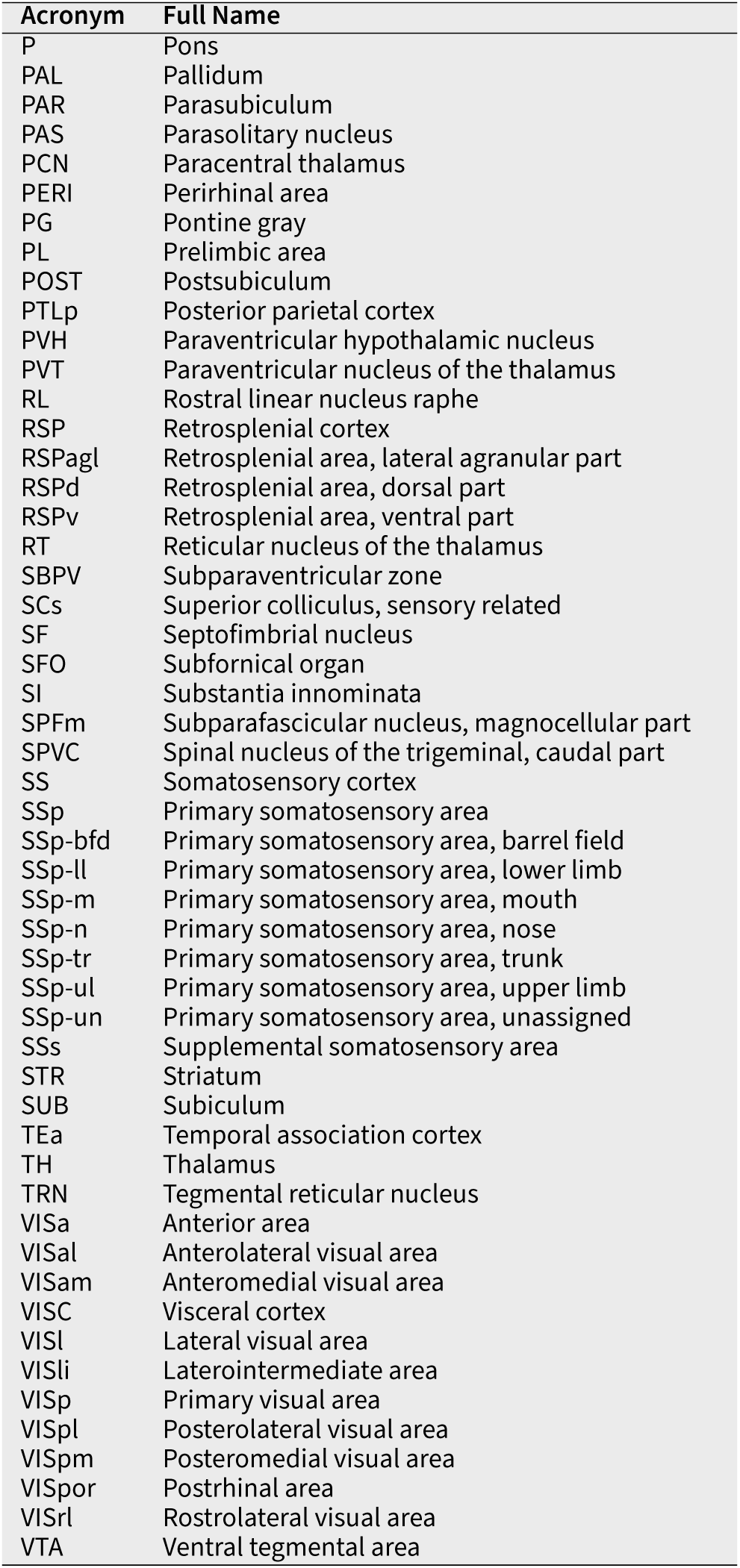
List of brain region acronyms and their full names.

**Table S5.** Neuroglancer links for Data in Figures.

| Figure | Purpose / Panels | Sample | Neuroglancer Link |
| --- | --- | --- | --- |
| Fig 5 | AiE2252m | 825027 | LINK |
| Fig 5 | Rorb-Cre | 780647 | LINK |
| Fig 5 | Ntsr1-Cre | 747936 | LINK |
| Fig 5 | Sert-Cre | 831959 | LINK |
| Fig 6 | LC (MOB) | 784354 | LINK |
| Fig 6 | LC (Spinal) | 762196 | LINK |
| Fig 6 | HIP | 797808 | LINK |
| Fig 7 | cFos | 751919 | LINK |
| Fig 7 | Ephys | 770311 | LINK |
| Fig 8 | ACA Inj 1 | 757189 (561) | LINK |
| Fig 8 | ACA Inj 2 | 757189 (488) | LINK |
| Fig 8 | ACA Inj 3 | 757188 | LINK |
| Fig 8 | ACA Inj 4 | 755809 | LINK |
| Fig 8 | ACA Inj 5 | 684814 | LINK |
| Fig 8 | ACA Inj 6 | 694512 | LINK |
| Fig 8 | MO Inj 1 | 684821 | LINK |
| Fig 8 | MO Inj 2 | 755808 (561) | LINK |
| Fig 8 | MO Inj 3 | 755808 (488) | LINK |
| Fig 8 | MO Inj 4 | 685904 | LINK |
| Fig 8 | MO Inj 5 | 761580 (488) | LINK |
| Fig 8 | MO Inj 6 | 761580 (561) | LINK |

## Notes

### Competing Interest Statement

The authors have declared no competing interest.

### Summary of Updates

One author was removed and we have written permission of this author in writing.

## References

1. A DSL for parallel and scalable computational pipelines| Nextflow [in en].

2. Achard, C., T. Kousi, M. Frey, M. Vidal, Y. Paychère, C. Hofmann, A. Iqbal, et al. 2024. “CellSeg3D: self-supervised 3D cell segmentation for light-sheet microscopy” [in en]. eLife 13.

3. Ahlers, J., D. Althviz Moré, O. Amsalem, A. Anderson, G. Bokota, P. Boone, J. Bragantini, et al. 2023. *Napari:* a multidimensional image viewer for Python.

4. Allen, W. E., L. A. DeNardo, M. Z. Chen, C. D. Liu, K. M. Loh, L. E. Fenno, C. Ramakrishnan, K. Deisseroth, & L. Luo. 2017. “Thirst-associated preoptic neurons encode an aversive motivational drive” [in eng]. Science 357: 1149–1155.

5. Allen Institute for Brain Science. 2026a. “Mouse Cardiac Perfusion Fixation and Brain Collection V.9” [in en].

6. Allen Institute for Brain Science. 2026b. “Retro-orbital Injection of AAV Vectors for Viral Genetic Tools Pipeline” [in en].

7. Allen Institute for Brain Science. 2026c. “Stereotaxic Injection by Nanoject Protocol V.8” [in en].

8. Amaya, A., C. Bennett, B. Ouellette, T. K. Ramirez, A. Williford, & R. Naidoo. 2026. “Whole Hemisphere Craniotomy for Electrophysiology” [in en].

9. Amaya, A., J. Swapp, A. Williford, & R. Howard. 2024. “General Setup and Takedown Procedures for Rodent Neurosurgery V.2” [in en].

10. anna.lakunina. 2024. “Duragel Application for Acute Electrophysiology Recordings” [in en]. ANTsX/ANTsPy. 2025.

11. Attarpour, A., J. Osmann, A. Rinaldi, T. Qi, N. Lal, S. Patel, M. Rozak, et al. 2025. “A deep learning pipeline for three-dimensional brain-wide mapping of local neuronal ensembles in teravoxel light-sheet microscopy” [in en]. Nature Methods, 1–12.

12. Avants, B., N. J. Tustison, & G. Song. 2009. “Advanced Normalization Tools: V1.0.” The Insight Journal.

13. Avants, B. B., N. J. Tustison, G. Song, P. A. Cook, A. Klein, & J. C. Gee. 2011. “A reproducible evaluation of ANTs similarity metric performance in brain image registration” [in eng]. NeuroImage 54: 2033–2044.

14. Bankhead, P., M. B. Loughrey, J. A. Fernández, Y. Dombrowski, D. G. McArt, P. D. Dunne, S. McQuaid, et al. 2017. “QuPath: Open source software for digital pathology image analysis” [in en]. Scientific Reports 7: 16878.

15. Becker, K., N. Jährling, S. Saghafi, R. Weiler, & H.-U. Dodt. 2012. “Chemical Clearing and Dehydration of GFP Expressing Mouse Brains” [in en]. PLOS ONE 7: e33916.

16. Ben-Simon, Y., M. Hooper, S. Narayan, T. L. Daigle, D. Dwivedi, S. W. Way, A. Oster, et al. 2025. “A suite of enhancer AAVs and transgenic mouse lines for genetic access to cortical cell types.” Cell 188: 3045–3064.e23.

17. Bennett, C., B. Ouellette, T. K. Ramirez, A. Cahoon, H. Cabasco, Y. Browning, A. Lakunina, et al. 2024. “SHIELD: Skull-shaped hemispheric implants enabling large-scale electrophysiology datasets in the mouse brain” [in en]. Neuron 112: 2869– 2885.e8.

18. Bria, A., & G. Iannello. 2012. “TeraStitcher - A tool for fast automatic 3D-stitching of teravoxel-sized microscopy images.” BMC Bioinformatics 13: 316.

19. Browning, Y., G. Lynch, C. King, A. Lakunina, M. Olsen, X. Yin, & J. Siegle. 2026. “Acute Neuropixels Recording Through a 3D Printed Implant” [in en].

20. Cheifet, B. 2021. “Promoting reproducibility with Code Ocean” [in eng]. Genome Biology 22: 65.

21. Chen, F., P. W. Tillberg, & E. S. Boyden. 2015. “Expansion microscopy.” Science 347: 543–548.

22. Chen, S., Y. Liu, Z. A. Wang, J. Colonell, L. D. Liu, H. Hou, N.-W. Tien, et al. 2024. “Brain-wide neural activity underlying memory-guided movement” [in English]. Cell 187: 676– 691.e16.

23. Chozinski, T. J., A. R. Halpern, H. Okawa, H.-J. Kim, G. J. Tremel, R. O. L. Wong, & J. C. Vaughan. 2016. “Expansion microscopy with conventional antibodies and fluorescent proteins” [in eng]. Nature Methods 13: 485–488.

24. Chung, K., & K. Deisseroth. 2013. “CLARITY for mapping the nervous system” [in en]. Nature Methods 10: 508–513.

25. Claudi, F., L. Petrucco, A. L. Tyson, T. Branco, T. W. Margrie, & R. Portugues. 2020. “BrainGlobe Atlas API: a common interface for neuroanatomical atlases” [in en]. Journal of Open Source Sokware 5: 2668.

26. Claudi, F., A. L. Tyson, L. Petrucco, T. W. Margrie, R. Portugues, & T. Branco. 2021. “Visualizing anatomically registered data with brainrender.” Edited by M. W. Mathis, K. M. Wassum, & J. Nunez-Iglesias. eLife 10: e65751.

27. Costantini, I., J.-P. Ghobril, A. P. Di Giovanna, A. L. Allegra Mascaro, L. Silvestri, M. C. Müllenbroich, L. Onofri, et al. 2015. “A versatile clearing agent for multi-modal brain imaging” [in eng]. Scientific Reports 5: 9808.

28. Dimidschstein, J., Q. Chen, R. Tremblay, S. L. Rogers, G.-A. Saldi, L. Guo, Q. Xu, et al. 2016. “A viral strategy for targeting and manipulating interneurons across vertebrate species” [in eng]. Nature Neuroscience 19: 1743–1749.

29. Dodt, H.-U., U. Leischner, A. Schierloh, N. Jährling, C. P. Mauch, K. Deininger, J. M. Deussing, et al. 2007. “Ultramicroscopy: three-dimensional visualization of neuronal networks in the whole mouse brain” [in en]. Nature Methods 4: 331–336.

30. Ertürk, A., K. Becker, N. Jährling, C. P. Mauch, C. D. Hojer, J. G. Egen, F. Hellal, et al. 2012. “Three-dimensional imaging of solvent-cleared organs using 3DISCO” [in eng]. Nature Protocols 7: 1983–1995.

31. Faress, I., V. Khalil, H. Yamamoto, S. Sajgo, K. Yonehara, & S. Nabavi. 2023. “Recombinase-independent AAV for anterograde transsynaptic tracing” [in eng]. Molecular Brain 16: 66.

32. Fischl, B. 2012. “FreeSurfer.” NeuroImage, 20 YEARS OF fMRI, 62: 774–781.

33. Friedmann, D., A. Pun, E. L. Adams, J. H. Lui, J. M. Kebschull, S. M. Grutzner, C. Castagnola, M. Tessier-Lavigne, & L. Luo. 2020. “Mapping mesoscale axonal projections in the mouse brain using a 3D convolutional network.” Proceedings of the National Academy of Sciences 117: 11068–11075.

34. Glaser, A., J. Chandrashekar, S. Vasquez, C. Arshadi, R. Javeri, N. Ouellette, X. Jiang, et al. 2025. “Expansion-assisted selective plane illumination microscopy for nanoscale imaging of centimeter-scale tissues” [in en]. eLife 12.

35. Goldman, D. B. 2010. “Vignette and Exposure Calibration and Compensation.” IEEE Transactions on Pattern Analysis and Machine Intelligence 32: 2276–2288.

36. Gong, S., M. Doughty, C. R. Harbaugh, A. Cummins, M. E. Hatten, N. Heintz, & C. R. Gerfen. 2007. “Targeting Cre recombinase to specific neuron populations with bacterial artificial chromosome constructs” [in eng]. The Journal of Neuroscience: The Official Journal of the Society for Neuroscience 27: 9817– 9823.

37. Gong, S., C. Zheng, M. L. Doughty, K. Losos, N. Didkovsky, U. B. Schambra, N. J. Nowak, et al. 2003. “A gene expression atlas of the central nervous system based on bacterial artificial chromosomes” [in en]. Nature 425: 917–925.

38. Graybuck, L. T., T. L. Daigle, A. E. Sedeño-Cortés, M. Walker, B. Kalmbach, G. H. Lenz, E. Morin, et al. 2021. “Enhancer viruses for combinatorial cell-subclass-specific labeling” [in eng]. Neuron 109: 1449–1464.e13.

39. Hama, H., H. Kurokawa, H. Kawano, R. Ando, T. Shimogori, H. Noda, K. Fukami, A. Sakaue-Sawano, & A. Miyawaki. 2011. “Scale: a chemical approach for fluorescence imaging and reconstruction of transparent mouse brain” [in en]. Nature Neuroscience 14: 1481–1488.

40. Harris, J. A., K. E. Hirokawa, S. A. Sorensen, H. Gu, M. Mills, L. L. Ng, P. Bohn, et al. 2014. “Anatomical characterization of Cre driver mice for neural circuit mapping and manipulation” [in eng]. Frontiers in Neural Circuits 8: 76.

41. He, K., X. Zhang, S. Ren, & J. Sun. 2015. Deep Residual Learning for Image Recognition.

42. Hörl, D., F. Rojas Rusak, F. Preusser, P. Tillberg, N. Randel, R. K. Chhetri, A. Cardona, et al. 2019. “BigStitcher: reconstructing high-resolution image datasets of cleared and expanded samples” [in en]. Nature Methods 16: 870–874.

43. Hörst, F., M. Rempe, L. Heine, C. Seibold, J. Keyl, G. Baldini, S. Ugurel, et al. 2024. “CellViT: Vision Transformers for precise cell segmentation and classification.” Medical Image Analysis 94: 103143.

44. Hou, B., D. Zhang, S. Zhao, M. Wei, Z. Yang, S. Wang, J. Wang, et al. 2015. “Scalable and DiI-compatible optical clearance of the mammalian brain” [in English]. Frontiers in Neuroanatomy 9.

45. Hrvatin, S., C. P. Tzeng, M. A. Nagy, H. Stroud, C. Koutsioumpa, O. F. Wilcox, E. G. Assad, et al. 2019. “A scalable platform for the development of cell-type-specific viral drivers.” Edited by A. E. West & C. Dulac. eLife 8: e48089.

46. Iqbal, A., A. Sheikh, & T. Karayannis. 2019. “DeNeRD: high-throughput detection of neurons for brain-wide analysis with deep learning” [in en]. Scientific Reports 9: 13828.

47. Jacques, S. L. 2013. “Optical properties of biological tissues: a review” [in eng]. Physics in Medicine and Biology 58: R37–61.

48. Johnson, G. A., Y. Tian, D. G. Ashbrook, G. P. Cofer, J. J. Cook, J. C. Gee, A. Hall, et al. 2023. “Merged magnetic resonance and light sheet microscopy of the whole mouse brain.” Proceedings of the National Academy of Sciences 120: e2218617120.

49. Ke, R., M. Mignardi, A. Pacureanu, J. Svedlund, J. Botling, C. Wählby, & M. Nilsson. 2013. “In situ sequencing for RNA analysis in preserved tissue and cells” [in eng]. Nature Methods 10: 857–860.

50. Kirst, C., S. Skriabine, A. Vieites-Prado, T. Topilko, P. Bertin, G. Gerschenfeld, F. Verny, et al. 2020. “Mapping the Fine-Scale Organization and Plasticity of the Brain Vasculature” [in English]. Cell 180: 780–795.e25.

51. Klein, S., M. Staring, K. Murphy, M. A. Viergever, & J. P. W. Pluim. 2010. “elastix: A Toolbox for Intensity-Based Medical Image Registration.” IEEE Transactions on Medical Imaging 29: 196– 205.

52. Krupa, O., G. Fragola, E. Hadden-Ford, J. T. Mory, T. Liu, Z. Humphrey, B. W. Rees, et al. 2021. “NuMorph: Tools for cortical cellular phenotyping in tissue-cleared whole-brain images.” Cell reports 37: 109802.

53. Ku, T., J. Swaney, J.-Y. Park, A. Albanese, E. Murray, J. H. Cho, Y.-G. Park, et al. 2016. “Multiplexed and scalable super-resolution imaging of three-dimensional protein localization in size-adjustable tissues” [in en]. Nature Biotechnology 34: 973–981.

54. Laboratory, I. B., K. Banga, J. Benson, J. Bhagat, D. Biderman, D. Birman, N. Bonacchi, et al. 2025. Reproducibility of in vivo electrophysiological measurements in mice [in en].

55. Lakunina, A., C. Grasso, B. Barad, & A. Amaya. 2025. “Dual Hemisphere Craniotomy for Electrophysiology” [in en].

56. Liang, X., Y. Zang, D. Dong, L. Zhang, M. Fang, X. Yang, A. Arranz, et al. 2016. “Stripe artifact elimination based on nonsubsampled contourlet transform for light sheet fluorescence microscopy” [in eng]. Journal of Biomedical Optics 21: 106005.

57. Liu, L. D., S. Chen, H. Hou, S. J. West, M. Faulkner, T. I. B. Laboratory, M. N. Economo, N. Li, & K. Svoboda. 2021. “Accurate Localization of Linear Probe Electrode Arrays across Multiple Brains” [in en]. eNeuro 8.

58. Luo, L., E. M. Callaway, & K. Svoboda. 2018. “Genetic Dissection of Neural Circuits: A Decade of Progress.” Neuron 98: 256– 281.

59. Madisen, L., A. R. Garner, D. Shimaoka, A. S. Chuong, N. C. Klapoetke, L. Li, A. van der Bourg, et al. 2015. “Transgenic mice for intersectional targeting of neural sensors and effectors with high specificity and performance” [in eng]. Neuron 85: 942–958.

60. Matho, K. S., D. Huilgol, W. Galbavy, M. He, G. Kim, X. An, J. Lu, et al. 2021. “Genetic dissection of the glutamatergic neuron system in cerebral cortex” [in eng]. Nature 598: 182–187.

61. Matsumoto, K., T. T. Mitani, S. A. Horiguchi, J. Kaneshiro, T. C. Murakami, T. Mano, H. Fujishima, et al. 2019. “Advanced CUBIC tissue clearing for whole-organ cell profiling” [in en]. Nature Protocols 14: 3506–3537.

62. Moore, J., D. Basurto-Lozada, S. Besson, J. Bogovic, J. Bragantini, E. M. Brown, J.-M. Burel, et al. 2023. “OME-Zarr: a cloud-optimized bioimaging file format with international community support” [in en]. Histochemistry and Cell Biology 160: 223–251.

63. Moritz, P., R. Nishihara, S. Wang, A. Tumanov, R. Liaw, E. Liang, M. Elibol, et al. 2018. Ray: a distributed framework for emerging AI applications. In Proceedings of the 13th USENIX conference on Operating Systems Design and Implementation, 561–577. OSDI’18. USA: USENIX Association. ISBN: 978-1-931971-47-8.

64. Murakami, T. C., T. Mano, S. Saikawa, S. A. Horiguchi, D. Shigeta, K. Baba, H. Sekiya, et al. 2018. “A three-dimensional single-cell-resolution whole-brain atlas using CUBIC-X expansion microscopy and tissue clearing” [in en]. Nature Neuroscience 21: 625–637.

65. Myers, H., & D. Toglia. 2025a. “Refractive Index Matching - EasyIndex” [in en].

65a. Myers, H., & D. Toglia. 2025b. “Whole Brain Embedding for SmartSPIM - EasyIndex with 2% Agarose” [in en].

67. Myers, H., & D. Toglia. 2025c. “Whole Mouse Brain Delipidation - LifeCanvas Active” [in en].

68. Narboux-Nême, N., L. M. Pavone, L. Avallone, X. Zhuang, & P. Gaspar. 2008. “Serotonin transporter transgenic (SERTcre) mouse line reveals developmental targets of serotonin specific reuptake inhibitors (SSRIs)” [in eng]. Neuropharmacology 55: 994–1005.

69. Neuroglancer — connectomics latest documentation.

70. Okuta, R., Y. Unno, D. Nishino, S. Hido, & Crissman. 2017. CuPy : A NumPy-Compatible Library for NVIDIA GPU Calculations.

71. Otsu, N. 1979. “A Threshold Selection Method from Gray-Level Histograms.” IEEE Transactions on Systems, Man, and Cybernetics 9: 62–66.

72. Peng, T., K. Thorn, T. Schroeder, L. Wang, F. J. Theis, C. Marr, & N. Navab. 2017. “A BaSiC tool for background and shading correction of optical microscopy images” [in en]. Nature Communications 8: 14836.

73. Perens, J., C. G. Salinas, J. L. Skytte, U. Roostalu, A. B. Dahl, T. B. Dyrby, F. Wichern, et al. 2021. “An Optimized Mouse Brain Atlas for Automated Mapping and Quantification of Neuronal Activity Using iDISCO+ and Light Sheet Fluorescence Microscopy” [in eng]. Neuroinformatics 19: 433–446.

74. Piccinini, F., E. Lucarelli, A. Gherardi, & A. Bevilacqua. 2012. “Multi-image based method to correct vignetting effect in light microscopy images” [in eng]. Journal of Microscopy 248: 6–22.

75. Pieper, S., M. Halle, & R. Kikinis. 2004. 3D Slicer. In 2004 2nd IEEE International Symposium on Biomedical Imaging: Nano to Macro (IEEE Cat No. 04EX821), 632–635 Vol. 1.

76. Pisano, T. J., A. T. Hoag, Z. M. Dhanerawala, S. R. Guariglia, C. Jung, H.-J. Boele, K. M. Seagraves, J. L. Verpeut, & S. S.-H. Wang. 2022. “Automated high-throughput mouse transsynaptic viral tracing using iDISCO+ tissue clearing, light-sheet microscopy, and BrainPipe.” STAR Protocols 3: 101289.

77. Qiu, S., Y. Hu, Y. Huang, T. Gao, X. Wang, D. Wang, B. Ren, et al. 2024. “Whole-brain spatial organization of hippocampal single-neuron projectomes” [in en]. Science 383: eadj9198.

78. Renier, N., E. L. Adams, C. Kirst, Z. Wu, R. Azevedo, J. Kohl, A. E. Autry, et al. 2016. “Mapping of brain activity by automated volume analysis of immediate early genes.” Cell 165: 1789– 1802.

79. Renier, N., Z. Wu, D. J. Simon, J. Yang, P. Ariel, & M. Tessier-Lavigne. 2014. “iDISCO: a simple, rapid method to immunolabel large tissue samples for volume imaging” [in eng]. Cell 159: 896–910.

80. Ricci, P., V. Gavryusev, C. Müllenbroich, L. Turrini, G. de Vito, L. Silvestri, G. Sancataldo, & F. S. Pavone. 2022. “Removing striping artifacts in light-sheet fluorescence microscopy: a review.” Progress in Biophysics and Molecular Biology, The Resolution Revolution: Fluorescence Microscopy of Biological Samples from Micro to Meso, 168: 52–65.

81. Rohde, J. 2023a. “Imaging cleared mouse brains on SmartSPIM” [in en].

80a. Rohde, J. 2023b. “SmartSPIM setup and alignment” [in en].

83. Rokicki, K., D. Schauder, D. J. Olbris, C. Goina, J. Clements, P. Edson, T. Kawase, et al. 2025.HortaCloud: An Open and Collaborative Platform for Whole Brain Neuronal Reconstructions [in en].

84. Roston, R. A., N. J. Tustison, & A. M. Maga. 2025. Anatomy-aware, label-informed approach improves image registration for challenging datasets [in en].

85. Schindelin, J., I. Arganda-Carreras, E. Frise, V. Kaynig, M. Longair, T. Pietzsch, S. Preibisch, et al. 2012. “Fiji: an open-source platform for biological-image analysis” [in en]. Nature Methods 9: 676–682.

86. Spalteholz, W. 1914. “Über das Durchsichtigmachen von menschlichen und tierischen Präparaten und seine theoretischen Bedingungen.” Verhandlungen der Anatomischen Gesellschak 28: 3–25.

87. Stelzer, E. H. K., F. Strobl, B.-J. Chang, F. Preusser, S. Preibisch, K. McDole, & R. Fiolka. 2021. “Light sheet fluorescence microscopy” [in en]. Nature Reviews Methods Primers 1: 73.

88. Stringer, C., T. Wang, M. Michaelos, & M. Pachitariu. 2021. “Cellpose: a generalist algorithm for cellular segmentation” [in en]. Nature Methods 18: 100–106.

89. Su, Z., P. Kosillo, K. Jung, S. Chen, M. T. Summers, A. Piet, H. Hou, et al. 2026. “Topographic structure and function of locus coeruleus norepinephrine neurons.” bioRxiv.

90. Sung, K., Y. Ding, J. Ma, H. Chen, V. Huang, M. Cheng, C. F. Yang, et al. 2016. “Simplified three-dimensional tissue clearing and incorporation of colorimetric phenotyping” [in en]. Scientific Reports 6: 30736.

91. Susaki, E. A., K. Tainaka, D. Perrin, F. Kishino, T. Tawara, T. M. Watanabe, C. Yokoyama, et al. 2014. “Whole-brain imaging with single-cell resolution using chemical cocktails and computational analysis” [in eng]. Cell 157: 726–739.

92. Tainaka, K., S. I. Kubota, T. Q. Suyama, E. A. Susaki, D. Perrin, M. Ukai-Tadenuma, H. Ukai, & H. R. Ueda. 2014. “Whole-Body Imaging with Single-Cell Resolution by Tissue Decolorization.” Cell 159: 911–924.

93. Tervo, D. G. R., B.-Y. Hwang, S. Viswanathan, T. Gaj, M. Lavzin, K. D. Ritola, S. Lindo, et al. 2016. “A Designer AAV Variant Permits Efficient Retrograde Access to Projection Neurons” [in English]. Neuron 92: 372–382.

94. Tillage, R. P., N. R. Sciolino, N. W. Plummer, D. Lustberg, L. C. Liles, M. Hsiang, J. M. Powell, et al. 2020. “Elimination of galanin synthesis in noradrenergic neurons reduces galanin in select brain areas and promotes active coping behaviors” [in eng]. Brain Structure & Function 225: 785–803.

95. Tomazevic, D., B. Likar, & F. Pernus. 2002. “Comparative evaluation of retrospective shading correction methods” [in eng]. Journal of Microscopy 208: 212–223.

96. Toyoshima, Y., T. Tokunaga, O. Hirose, M. Kanamori, T. Teramoto, M. S. Jang, S. Kuge, et al. 2016. “Accurate Automatic Detection of Densely Distributed Cell Nuclei in 3D Space” [in en]. PLOS Computational Biology 12: e1004970.

97. Turner, M. A., T. Chartrand, M. T. Summers, M. Hooper, C. v. Velthoven, J. Waters, S. d. Vries, et al. 2025. Exploring the correspondence between gene expression and thalamic nuclei using the THALMANAC resource [in en].

98. Tustison, N. J., B. B. Avants, P. A. Cook, Y. Zheng, A. Egan, P. A. Yushkevich, & J. C. Gee. 2010. “N4ITK: Improved N3 Bias Correction.” IEEE transactions on medical imaging 29: 1310– 1320.

99. Tustison, N. J., P. A. Cook, A. J. Holbrook, H. J. Johnson, J. Muschelli, G. A. Devenyi, J. T. Duda, et al. 2021. “The ANTsX ecosystem for quantitative biological and medical imaging” [in en]. Scientific Reports 11: 9068.

100. Tyson, A. L., C. V. Rousseau, C. J. Niedworok, S. Keshavarzi, C. Tsitoura, L. Cossell, M. Strom, & T. W. Margrie. 2021a. “A deep learning algorithm for 3D cell detection in whole mouse brain image datasets” [in en]. PLOS Computational Biology 17: e1009074.

101. Tyson, A. L., C. V. Rousseau, C. J. Niedworok, S. Keshavarzi, C. Tsitoura, L. Cossell, M. Strom, & T. W. Margrie. 2021b. “A deep learning algorithm for 3D cell detection in whole mouse brain image datasets” [in en]. PLOS Computational Biology 17: e1009074.

102. Tyson, A. L., M. Vélez-Fort, C. V. Rousseau, L. Cossell, C. Tsitoura, S. C. Lenzi, H. A. Obenhaus, et al. 2022. “Accurate determination of marker location within whole-brain microscopy images” [in en]. Scientific Reports 12: 867.

103. Ueta, Y., H. Yamashita, M. Kawata, & K. Koizumi. 1995. “Water deprivation induces regional expression of c-*fos* protein in the brain of inbred polydipsic mice.” Brain Research 677: 221–228.

104. Varga, D. 2020. “No-Reference Image Quality Assessment Based on the Fusion of Statistical and Perceptual Features.” Journal of Imaging 6: 75.

105. Wang, Q., S.-L. Ding, Y. Li, J. Royall, D. Feng, P. Lesnar, N. Graddis, et al. 2020. “The Allen Mouse Brain Common Coordinate Framework: A 3D Reference Atlas.” Cell 181: 936–953.e20.

106. Wang, L.-W., Y.-L. Wu, C.-L. Lee, C.-C. Cheng, K.-Y. Lu, J.-H. Tsai, Y.-H. Lin, et al. 2023. A Weakly Supervised U-Net Model for Precise Whole Brain Immunolabeled Cell Detection [in en].

107. Wilkinson, M. D., M. Dumontier, I. J. Aalbersberg, G. Appleton, M. Axton, A. Baak, N. Blomberg, et al. 2016. “The FAIR Guiding Principles for scientific data management and stewardship” [in en]. Scientific Data 3: 160018.

108. Yamawaki, N., K. Borges, B. A. Suter, K. D. Harris, & G. M. G. Shepherd. 2014. A genuine layer 4 in motor cortex with prototypical synaptic circuit connectivity [in en].

109. Yang, B., J. B. Treweek, R. P. Kulkarni, B. E. Deverman, C.-K. Chen, E. Lubeck, S. Shah, L. Cai, & V. Gradinaru. 2014. “Single-cell phenotyping within transparent intact tissue through whole-body clearing” [in eng]. Cell 158: 945–958.

110. Yao, Z., C. T. J. van Velthoven, M. Kunst, M. Zhang, D. McMillen, C. Lee, W. Jung, et al. 2023. “A high-resolution transcriptomic and spatial atlas of cell types in the whole mouse brain” [in en]. Nature 624: 317–332.

111. Yin, X., anna.lakunina, & J. Siegle. 2024. “Preparing a 3D Printed Implant for Acute In Vivo Electrophysiology V.1” [in en].

112. Zaharia, M., M. Chowdhury, M. J. Franklin, S. Shenker, & I. Stoica. 2010. Spark: cluster computing with working sets. In Proceedings of the 2nd USENIX conference on Hot topics in cloud computing, 10. HotCloud’10. USA: USENIX Association.

113. Zheng, W., H. Mu, Z. Chen, J. Liu, D. Xia, Y. Cheng, Q. Jing, et al. 2024. “NEATmap: a high-efficiency deep learning approach for whole mouse brain neuronal activity trace mapping.” National Science Review 11: nwae109.

114. Zhuang, X., J. Masson, J. A. Gingrich, S. Rayport, & R. Hen. 2005. “Targeted gene expression in dopamine and serotonin neurons of the mouse brain” [in eng]. Journal of Neuroscience Methods 143: 27–32.

115. Zingg, B., X.-L. Chou, Z.-G. Zhang, L. Mesik, F. Liang, H. W. Tao, & L. I. Zhang. 2017. “AAV-Mediated Anterograde Transsynaptic Tagging: Mapping Corticocollicular Input-Defined Neural Pathways for Defense Behaviors” [in eng]. Neuron 93: 33–47.

116. Zingg, B., B. Peng, J. Huang, H. W. Tao, & L. I. Zhang. 2020. “Synaptic Specificity and Application of Anterograde Transsynaptic AAV for Probing Neural Circuitry” [in eng]. The Journal of Neuroscience: The Official Journal of the Society for Neuroscience 40: 3250–3267.

